# A pennycress *transparent testa 8* knockout mutant has drastic changes in seed coat anatomy and chemical compositions

**DOI:** 10.1101/2025.03.10.642428

**Authors:** Xinxin Ding, Summer Duckworth, Madeline Southworth, Barsanti Gautam, Andrew Lipton, Dusan Velickovic, John C. Sedbrook, Pubudu Handakumbura

## Abstract

Pennycress is a winter annual intermediate crop with approximately 30% seed oil content suitable for producing biofuels. Here, we evaluated seed development, anatomy, and agronomically relevant traits of a *transparent testa 8* knockout mutant (*tt8-2bp*) generated by CRISPR genome editing to improve seed quality. We performed histochemical analyses on wild-type and *tt8-2bp* seeds at different developmental stages. No visible anatomical defects were observed in embryos and endosperm of *tt8-2bp* seeds. However, *tt8-2bp* seed coats completely lost proanthocyanidins which were accumulated in an inner integument cell layer and in the thickened cell wall of an outer integument cell layer of wild-type seed coats. Based on spatial metabolomic and solid-state NMR analyses, *tt8-2bp* seed coats had decreased aromatic compounds and cell wall polysaccharides compared to wild-type seed coats. Additionally, *tt8-2bp* seeds had reduced seed coat dry weights and increased embryo dry weights compared to wild-type seeds, indicating changes in macronutrient partitioning during seed development. Mature *tt8-2bp* seeds exhibited increased imbibition rates and seed coat permeability to water-soluble molecules, suggesting a higher seed coat hydrophilicity than wild-type seeds. In conclusion, we did not find defects in *tt8-2bp* mutant seeds that were unfavorable agronomically, supporting that *TT8* is an attractive target for pennycress domestication.

**Highlight:** Histochemical analyses of pennycress seeds revealed a complete loss of proanthocyanidins in *tt8-2bp* seed coats accompanied by increased seed imbibition rates and seed coat permeability compared to wild-type seeds.

## Introduction

Pennycress (*Thlaspi arvense* L.) is an emerging intermediate oilseed crop in the *Brassicaceae* family with many advantages. Pennycress is diploid and has a draft genome assembled (Dorn *et al*., 2015), and its protein coding genes are highly similar to those of Arabidopsis (*Arabidopsis thaliana*). A comparative analysis using 65% protein sequence identity cutoff identified 77.5% of annotated pennycress genes with at least one Arabidopsis ortholog and 52.8% pennycress genes have one-to-one correspondence with their Arabidopsis orthologs (Chopra *et al*., 2018). Additionally, pennycress can be transformed *via Agrobacterium*-mediated vacuum infiltration with 0.5% transformation efficiency, enabling genetic modification and genome editing (McGinn *et al*., 2019). The high similarity of protein encoding genes between pennycress and Arabidopsis and the availability of molecular tools help accelerate pennycress domestication *via* translational research. Pennycress has very high cold tolerance and a short life cycle which makes it suitable for planting between corn and soybean growth seasons (Lee *et al*., 2024). Pennycress seeds have high seed oil contents ranging between 29-32% on a seed dry weight basis (Moser *et al*., 2009; Chopra *et al*., 2019), and the seed oil profile allows for easy conversion to biofuels, which can help fulfill demands for sustainable aviation fuel (Moser *et al*., 2009; Phippen *et al*., 2022). Moreover, pennycress seeds have high protein contents ranging between 33-46% on a seed dry weight basis (Tsogtbaatar *et al*., 2015; Arias *et al*., 2023). To improve pennycress seed meal for animal feed, the seed glucosinolate and erucic acid contents must be reduced by traditional and/or mutation breeding due to their negative effects on animal health (Tripathi and Mishra, 2017; Lee *et al*., 2024).

Improving pennycress seed quality and the total seed oil content is central to pennycress domestication (Chopra *et al*., 2020; Esfahanian *et al*., 2021; Jarvis *et al*., 2021; Hartnell *et al*., 2023; Guzha *et al*., 2024). Seed quality, especially seed germination and vigor, determines the speed and uniformity of seedling establishment in the field, which directly affects crop density, leaf canopy establishment, and crop yields (Kumar SP *et al*., 2016; Reed *et al*., 2022). Seed dormancy is prevalent in pennycress seeds, especially when freshly harvested (Hazebroek and Metzger, 1990; Chen *et al*., 2022). Dormancy can be broken by treating the seeds with gibberellic acid or storing the seeds at room temperature for several months (Sedbrook *et al*., 2014; Koirala *et al*., 2022), but these methods increase the seed processing cost and time cost before planting. Thus, reduced seed dormancy is a critical trait for pennycress domestication.

Additionally. increasing total seed oil content is crucial for improving the economic value of oilseed crops. A soybean study estimated that 20 million dollars would be lost in revenue with every 1% reduction in seed oil content regardless of yields due to increased costs of processing more seeds to obtain the same amount of oil (Digrado *et al*., 2024). Many studies have been done on *Brassica napus* seeds (*e.g.*, rapeseed and canola) to investigate if there are associations among seed dormancy, germination time, oil contents, and fatty acid compositions in different cultivars (Gu *et al*., 2019; Brown *et al*., 2023; Brown *et al*., 2024). Overall, only weak correlations were found between seed germination rates and seed oil contents or fatty acid compositions (Gu *et al*., 2019; Brown *et al*., 2023; Brown *et al*., 2024). This suggests that it is possible to simultaneously improve seed germination rates and oil contents.

Arabidopsis studies show that knockout (KO) mutations of several *TRANSPARENT TESTA* (*TT*) genes, such as *TTG1* (Chen *et al*., 2015a), *TT2* (Chen *et al*., 2012), and *TT8* (Chen *et al*., 2014), can simultaneously increase total seed oil contents and reduce seed dormancy (Debeaujon *et al*., 2000). In Arabidopsis, *TTG1*, *TT2*, and *TT8* encode transcription factors which form a transcription factor complex and synergistically regulate the flavonoid biosynthesis pathway (Baudry *et al*., 2004). The TTG1-TT2-TT8 transcription factor complex activates anthocyanidin reductase encoded by *BANYULS*, a core enzyme in proanthocyanidin (PA) biosynthesis (Debeaujon *et al*., 2003). In Arabidopsis, *BANYULS* is activated specifically in the seed coat inner integument and the chalazal where PAs accumulate (Debeaujon *et al*., 2003). PAs are colorless flavonoid polymers which turn brown after polymerization into condensed tannins by oxidation (Xie *et al*., 2003; Feng *et al*., 2014). KO mutations in the pennycress orthologs of *TTG1*, *TT2*, and *TT8* also generated seeds with transparent testa, demonstrating that the function of the TTG1-TT2-TT8 transcription factor complex is likely conserved in pennycress (Chopra *et al*., 2018). Consistently, pennycress *tt2* KO mutant seeds have reduced dormancy and high germination rates (>89%) similar to Arabidopsis *tt2* KO mutant seeds (Debeaujon *et al*., 2000; Ott *et al*., 2021). Additionally, pennycress *tt8* KO mutant seeds have reduced dormancy and fiber contents and increased oil and protein contents compared to wild-type seeds (Sedbrook and Durrett, 2020; Koirala *et al*., 2023). These results suggest that pennycress *TT2* and *TT8* are good targets for improving pennycress seed quality and nutrient profile. However, there has not been any study systematically characterizing the seed development and anatomy of pennycress seeds, which is essential for understanding the cellular and molecular mechanisms underlying development of different seed tissues. Additionally, we need to evaluate the effects of *tt2* and *tt8* KO mutations on seed development, seed chemical compositions, and other agronomically relevant seed traits to understand what impacts these mutations will have on pennycress domestication.

Seed development and anatomy have been widely studied in Arabidopsis and were used as references for this study. In angiosperms, the embryo and endosperm derive from a double fertilization event while the seed coat develops from the integuments of the ovule (Lafon-Placette and Köhler, 2014; Huang *et al*., 2023). After fertilization, the three main seed tissues follow very different developmental programs. There are two main phases in embryo development, *i.e.*, embryo morphogenesis and maturation. In the first phase, the apical–basal pattern of the basic embryo body is established, and in the second phase the embryo accumulates storage compounds and undergoes dehydration (Jürgens, 2001; Huang *et al*., 2023). The endosperm development also has two main phases, *i.e.*, the syncytial phase and cellular phase, where the endosperm initially undergoes mitosis without cell wall formation and the cellularization occurs later (Lafon-Placette and Köhler, 2014; Huang *et al*., 2023). During embryo development, endosperm cells gradually undergo programmed cell death, and only the periphery endosperm layer stays alive at seed maturity (Huang *et al*., 2023). In contrast to embryo and endosperm, seed coat develops solely from maternal tissues. In the ovule, the integument and nucellus enclose the embryo sac. The number of integument layers varies among different species, but most angiosperms have two integuments named inner and outer integuments (Huang *et al*., 2023). In Arabidopsis, the inner integument (ii) has three cell layers, ii1, ii1’, and ii2, and the outer integument (oi) has two cell layers, oi1 and oi2 (Huang *et al*., 2023). The ii1 layer of Arabidopsis seed coat synthesizes and accumulates PAs, conferring a brown color and increased mechanical strength as well as modulating seed coat permeability in mature seeds (Huang *et al*., 2023).

In this study, we focused on a pennycress *tt8* KO mutant, *tt8-2bp*, which was generated by clustered regularly interspaced short palindromic repeats (CRISPR) gene editing, targeting a two base-pair deletion within the *TT8* coding sequence. The *tt8-2bp* mutation was introduced into the pennycress Spring32-10 (Spring32) background which is an inbred line generated for basic research purposes. Spring-type pennycress seeds including Spring 32 have little to no dormancy and the plants can flower without vernalization (McGinn *et al*., 2019). Comparing Spring32 wild type to the *tt8-2bp* mutant, we investigated the effects of *tt8* KO mutations on pennycress seed development and anatomy, chemical compositions of different seed tissues, and agronomically relevant seed traits including seed coat permeability, seed imbibition rates, dry weights of seed coats and embryos, and seed coat gas permeability. To characterize the anatomy of developing wild-type and *tt8-2bp* seeds, we performed histological staining on seeds at 7, 11, 15, 19, 23, and 27 days after pollination (DAP). Based on metabolite profiles of developing pennycress seeds (Tsogtbaatar *et al*., 2015), pennycress embryos transition from morphogenesis to maturation between 7-27 DAP, which would enable us to characterize major developmental changes in embryo, endosperm, and seed coat. To analyze changes in seed chemical compositions, we performed spatial metabolomic profiling on 27 DAP seeds and solid-state nuclear magnetic resonance (ss NMR) analysis on mature seed coats of wild type and *tt8-2bp*. We did not find any anatomical defects in the embryo and endosperm of *tt8-2bp* seeds but did detect drastic changes in seed coat anatomy and chemical compositions of the embryo and seed coat of *tt8-2bp* compared to wild-type seeds. The changes in anatomy and chemical compositions of *tt8-2bp* seed coats were correlated with significantly decreased seed coat dry weights and increased embryo dry weights, seed coat permeability to water-soluble molecules, and seed imbibition rates. The detailed development map of Spring32 wild-type and *tt8-2bp* seeds at 7-27 DAP provides important information for understanding cellular and molecular mechanisms of pennycress seed tissue development and grain filling. The PA deficiency and reduced dry weights of seed coats and increased small sugars, fatty acids, and embryo dry weights of *tt8-2bp* seeds suggest pennycress *TT8* expression modulates nutrient partitioning between pennycress seed coats and embryos.

## Materials and Methods

### Plant growth

The *tt8-2bp* KO allele was generated by Maliheh Esfahanian and consists of a 2 base-pair (bp) deletion located 179 bp downstream of the translational start site resulting in a frameshift mutation. Mature seeds of Spring32-10 wild-type (hereafter referred to as Spring32) and *tt8-2bp* were first germinated in petri dishes before transferred into pots. For sterilization, the seeds were transferred into 2 mL Eppendorf tubes and rinsed with 70% ethanol three times by adding 1 mL of 70% ethanol, quickly vortexing the tubes for 3 seconds, and disposing the ethanol each time. After sterilization, the seeds were rinsed with sterile milli-Q water three times following the same steps as before. Then, the seeds were placed on a piece of Whatman paper autoclaved before use in a 100 x 15 mm sterile plastic petri dish where 2 mL of sterile milli-Q water were added, and the petri dishes were sealed with parafilm. The petri dishes were placed in the growth chamber where the plants were later grown for 2-4 days. After most seeds were germinated, the young seedlings were transferred into pots (3.5" W x 3.5" L x 3.5" D) with pre-wetted commercial growth medium (PRO-MIX® BX Mycorrhizae™ Growing Medium). All the pots were kept in trays (10.94" W x 21.44" L x 2.44" D, 18 pots in a full tray) and the trays were covered with transparent plastic domes for two days to help keep the pots moist while the seedlings establishing their root system. All the plants were grown in a growth chamber under a 16h/8h light/dark cycle at 22 °C and 50% humidity under a constant light intensity of 200 μmol m^−2^ s^−1^ during daytime and at 20 °C and 70% humidity during nighttime. Watering and fertilization were done by adding deionized (DI) water and liquid fertilizer (15-16-17 Peat-Lite fertilizer, Jack’s Professional, product #77220), respectively, directly into the tray. The liquid fertilizer was made to have 100 ppm N feed rate (*i.e.*, 0.67g solid fertilizer per liter of DI water).

Before the emergence of the third pairs of true leaves, 1 L DI water was added to each tray on Tuesday and Friday of each week. After the emergence of the third pairs of true leaves, 1 L of fertilizer was also added to each tray on Friday each week in addition to DI water. After bolting, 1 L of fertilizer was also added to each tray on Tuesday and Friday each week in addition to DI water. Once seed pods are formed, 2 L DI water and 1 L fertilizer were added to each tray on Tuesday and Friday each week until seed harvest or all the seed pods were senesced.

### Manual pollination and chemical fixation of pennycress seeds

To obtain seeds at different developmental stages, flowers were manually pollinated and the seeds were harvested on different days after pollination to be preserved for histology and MALDI analyses. To ensure that after manual pollination all the seeds developed under a comparable condition, each plant was only pollinated once at 4-8 days after the opening of the first flower. Based on earlier studies on pennycress flowers and seeds (Tsogtbaatar *et al*., 2015; Thomas *et al*., 2017), it was shown that pennycress flowers are larger than Arabidopsis flowers but have very similar flower structures, and thus we can pollinate pennycress manually in the same way as done with Arabidopsis flowers. Because pennycress can produce multiple flower stems, we only pollinated flowers from the lead flower stem at 6-10 days after we observed the first flower opening on the lead stem to ensure consistency in the seed samples we harvested. For histological analysis, seed pods from manually pollinated flowers were harvested at specific developmental stages, and the seeds were fixed in fixation solution consisted of formalin:ethanol:acetic acid:water (10:50:5:35 v/v/v/v) with 4% formaldehyde in glass vials. To ensure quick fixation, vacuum infiltration was applied for 15 minutes after soaking the seeds in the fixation solution. The glass vials were then sealed and stored at 4 °C until analysis.

### Paraffin embedding of fixed pennycress seeds and histological analyses

Chemically fixed seeds were transferred from the fixation solution into a petri dish with 60% ethanol. Five tiny punctures were made on each side of the seeds to help melted paraffin diffuse inside the seeds in later steps. The punctures were crucial since intact seed coats prevented melted paraffin diffuse inside. Then, the seeds were dehydrated through a graded ethanol series, cleared in xylenes, and embedded in paraffin wax (**Supplementary Tab. S1**). Sections (8-μm thickness) were cut using a Mcrom Heidelberg rotary microtome (Model HM330) and transferred onto glass microscope slides with frosted ends. The slides were labeled with pencils, so the writings do not fade in the deparaffinization and staining process later. The slides were then incubated at 57 °C for 1hr to make the sections adhere tightly to the slide.

Before staining, the sections were first deparaffinized with xylene and then rehydrated by incubating in a decreasing alcoholic series (**Supplementary Tab. S2**). The alcohol concentration in the last step of rehydration should be the same as the alcohol concentration of the staining solution used immediately after. For unstained control, the sections were rehydrated to DI water, then dehydrated in 100% ethanol and xylenes in consecutive steps for one minute each step, and mounted with Permount mounting medium (Fisher Chemical) before covering the sections with a coverslip. All the unstained and stained samples were visualized with either 10x, 20x, or 40x objectives (Zeiss Fluar 10x/0.50, Zeiss LD Plan-Neofluar 20x/0.4 Corr M27, and Zeiss LD Plan-Neofluar 40x/0.60 Corr) and images were captured with a Zeiss Axiocam 305 color camera on a Zeiss PALM Microbeam Laser Capture Microdissection System. The staining conditions using different dyes are described below:

The seed paraffin sections were stained with alcian blue and safranin O to visualize primary and secondary cell wall. After rehydration to 50% ethanol, the sections were first stained for one minute in 1.5% Safranin O (w/v) dissolved in 50% ethanol followed by incubating the slides with DI water for four times, 20 seconds each time. Then, the sections were stained with 1% alcian blue (3% acetic acid, pH 2.5) for three minutes followed by incubating the slides in DI water for one minute. Finally, the sections were differentiated in 95% ethanol for 1.5 minutes, dehydrated in 100% ethanol and then xylenes for one minute each, and mounted with Permount mounting medium (Fisher Chemical) before covering the sections with a coverslip.

The seed paraffin sections were stained with alcian blue and Lugol’s iodine to visualize primary cell wall components and starch granules. This staining protocol was adapted from (Kutík and Beneš, 1977). The sections were rehydrated to 50% ethanol and then incubated in DI water for three minutes. The sections were first stained with 1% alcian blue (3% acetic acid, pH 2.5), incubated in DI water, and differentiated in 95% ethanol as described earlier. Then, the sections were incubated in DI water for one minute and stained in 0.25% Lugol’s iodine (VIG-271, Volu-Sol) diluted in DI water for seven minutes followed by incubating the sections twice with DI water, 20 seconds each time. Finally, the sections were mounted with a pre-made mounting medium (40% Arabic gum, 0.05% Lugol’s iodine, 59.95% DI water, w/w/v) to prevent the color fading of stained starch granules. The mounting medium can be stored at 4 °C in dark for one week. After mounting and covering the sections with a coverslip, the slides were visualized immediately for this study. Alternatively, the mounted slides can be kept in dark to dry overnight before visualization.

The seed paraffin sections were stained with phloroglucinol-hydrochloric acid (HCl), *i.e.*, Wiesner staining solution, to visualize lignin in cell wall. Sections were rehydrated to 70% ethanol. One to two drops of freshly made phloroglucinol-HCl staining solution (2% phloroglucinol, 12% HCl, 21% water, 65% ethanol, w/v/v/v) were added on the top of the sections (Pradhan Mitra and Loqué, 2014). The sections were covered with a coverslip before visualization. The staining solution dried up in 10 minutes, so the visualization was done within this time frame.

The seed paraffin sections were stained with freshly made vanillin-hydrochloric acid (HCl) solution to visualize proanthocyanidins. This staining protocol was adapted from (Feng *et al*., 2014; Xuan *et al*., 2014). Sections were incubated in 90% methanol for five minutes after deparaffinized with xylenes. Then, two drops of freshly made vanillin-HCl solution (1% vanillin, 90% methanol, 3.7% HCl, 5.3% DI water, w/v/v/v) was added on top of the sections. The sections were covered with a coverslip before visualization. The staining solution dried up in 10 minutes, so the visualization was done within this time frame.

The seed paraffin sections were stained with 0.05% (w/v) of ruthenium red solution to detect pectin in cell wall. The sections were rehydrated to 50% ethanol and then incubated in DI water for five minutes. Then, the sections were stained with 0.05% (w/v) of ruthenium red (11103-72-3, MilliporeSigma) dissolved in DI water followed by rinsing the sections twice with DI water, 20 seconds each time. Finally, the slides were dehydrated in 100% ethanol and xylenes for one minute each, and mounted with Permount mounting medium (SP15-100, Fisher Chemical) before covering the sections with a coverslip.

### Cryosectioning and staining of 15 DAP pennycress seeds to detect PAs

Pennycress wild-type and *tt8-2bp* seeds at 15 DAP were flash frozen immediately after harvest and stored at −80 °C until analysis. The seeds were embedded in a viscous embedding solution (7.5% hydroxypropyl methylcellulose [HPMC], 2.5% polyvinylpyrrolidone [PVP], 90% Milli-Q water, w/w/v). The embedding solution was pre-chilled on ice before use to prevent the seeds from thawing during the embedding process. First, a thin layer of embedding solution was poured into an embedding mold and chill on dry ice. Before the embedding solution was completely frozen, quickly put pennycress seeds in the desired location and orientation on top of the thin layer of embedding solution. Finally, pour more embedding solution to completely cover the seeds and let the solution freeze on dry ice. Store the embedded seeds at −80 °C until sectioning. The embedded seeds were sectioned at −12°C with a crytome (CryoStar NX-70 Cryostat, Thermo Scientific, Runcorn, UK), and 25-μm sections were thaw mounted on glass microscope slides. The slides were vacuum dried in a desiccator immediately after sectioning. To visualize proanthocyanidins, two drops of freshly made vanillin-HCl solution (1% vanillin, 90% methanol, 3.7% HCl, 5.3% DI water, w/v/v/v) was added on top of the sections. For unstained control sections, two drops of HCl solution (90% methanol, 3.7% HCl, 5.3% DI water, v/v/v) was added on top of the sections. The sections were covered with a coverslip before visualization.

### Mature seed mucilage staining

Mature pennycress wild-type and *tt8-2bp* seeds were stained with ruthenium red based on a published protocol for staining Arabidopsis mature seeds (McFarlane *et al*., 2014). To verify that the staining was done properly, mature Arabidopsis Col-0 seeds were also stained at the same time. Briefly, 0.01% ruthenium red solution (Sigma-Aldrich, catalog number: 11103-72-3) and 50 mM EDTA (pH 7.5) were prepared on the day of staining. The seeds were placed in 2 mL Eppendorf tubes and 800 μL of 50 mM EDTA (pH 7.5) was added. The tubes were incubated on an orbital shaker (400 rpm) for two hours at room temperature to promote the release of mucilage from the outer seed coat. Then, the 50 mM EDTA was removed from all tubes and replaced with 800 μL of 0.01% ruthenium red solution. The tubes were incubated for another hour under the same condition as before. Finally, the ruthenium red solution was removed and replaced with Milli-Q water. Immediately, the seeds were observed under a dissecting microscope.

### Seed MALDI-MSI analysis

Pennycress wild-type and *tt8-2bp* seeds at 27 DAP were flash frozen immediately after harvest and stored at −80 °C until analysis. The seeds were embedded as described under “Cryosectioning and staining of 15 DAP pennycress seeds to detect PAs”. The cryosectioning, sample preparation, and MALDI-MSI data collection protocols were adapted from (Balasubramanian *et al*., 2024) with a few adjustments. For all the seed samples, 20-μm sections were thaw mounted on indium tin oxide (ITO)-coated glass slides. For both wild-type and *tt8-2bp* seeds, cryosections were mounted on two different slides to be later analyzed under positive and negative ion mode, respectively. The sections were vacuum dried and coated with 2,5-DHB (2,5-dihydroxybenzoic acid) and NEDC (N-naphthylethylenediamine dihydrochloride) MALDI matrices using M5-Sprayer (HTX Technologies, Chapel Hill, NC) for analysis under positive and negative ion mode, respectively, as described in (Balasubramanian *et al*., 2024). DHB was prepared at a concentration of 40 mg/mL DHB (in 70% MeOH) and was sprayed at 50 µL/min flow rate using the same M5-Sprayer. The nozzle temperature was set to 70 °C, with 12 cycles at 3 mm track spacing with a crisscross pattern. A 2 s drying period was added between cycles, a linear flow was set to 1,200 mm/min with 10 PSI of nitrogen gas and a 40 mm nozzle height. NEDC was prepared at a concentration of 7 mg/mL NEDC in 70% MeOH and was sprayed at 120 µL/min flow rate using the same M5-Sprayer. The nozzle temperature was set to 70 °C, with 8 cycles at 3 mm track spacing with a crisscross pattern. A 0 s drying period was added between cycles, a linear flow was set to 1200 mm/min with 10 PSI of nitrogen gas and a 40 mm nozzle height. All imaging was performed on a Bruker Daltonics 12T solariX FTICR MS, equipped with a ParaCell. This instrument has an Apollo II ESI and MALDI source with a SmartBeam II frequency-tripled (355 nm) Nd: YAG laser (Bremen, Germany). For both positive and negative ion mode analyses, data acquisition was done using with broadband excitations from *m/z* 100 to 500 for detecting small molecules and from *m/z* 500 to 2,000 for detecting lipids and larger molecules. The observed mass resolution was 110k at *m/z* 400. The ion images were generated *via* FlexImaging (Bruker Daltonics, v.5.0) with a 25-μm step size (*i.e.*, a 25 x 25-μm^2^ spatial resolution. The generated ion images were annotated automatedly based on the centroided dataset using the METASPACE platform, which is a spatial metabolomics knowledge database (Ovchinnikova *et al*., 2020). For the data analysis of this study, KEGG-v1 and SwissLipids were used to annotate the small molecule and lipid datasets, respectively, within 3 ppm m/z tolerance.

For all the relative quantitative analyses comparing metabolite abundances in wild-type vs. *tt8* seeds, at least three biological replicates, *i.e.*, MALDI-MSI data collected from sections of three different seeds, were used. All the small molecules and lipids analyzed in this study were based on data collected under the negative ion mode. The whole seed and embryo regions were manually selected in the METASPACE based on the brightfield images of all seed sections, and the ion intensities of all the selected regions were exported in a table for all the small molecules and lipids of interest. For the selected whole seed and embryo regions, the total ion intensities and total number of pixels were calculated from the exported tables. For the seed coat region, the total ion intensities and number of pixels were calculated by subtracting the values of the embryo from those of the whole seed. Depending on the ion signal localization patterns of different small molecules and lipids, the average ion intensity per pixel was calculated by dividing the total ion intensities of a whole seed section by the total number of pixels of the whole seed, seed coat, or embryo regions. The average ion intensities of proanthocyanidin A2 (C_30_H_24_O_12_), many aromatic compounds (C_15_H_12_O_6_), digalacturonate and glycyrrhizin (C_12_H_18_O_13_), and some sugar acids and sugar acid derivatives (C_6_H_10_O_7_) were calculated with the total pixel numbers of the seed coat regions. The average ion intensity of sinigrin (C_10_H_17_NO_9_S_2_) was calculated with the total pixel numbers of the embryo regions. The average ion intensities of the rest of the molecular features of interest were calculated with the total pixel numbers of the whole seeds for statistical tests.

### Ss NMR analysis of the mature seed outer seed coats

Mature pennycress wild-type and *tt8-2bp* seeds, respectively, were pooled from three different plants of each genotype. From each plant, 30 seeds were randomly selected from among all the seeds harvested, and 90 seeds were obtained for each genotype. The seeds were put in 2 mL Eppendorf tubes and rinsed three times with 70% ethanol to remove any foreign substances. Then, the seeds were rinsed three times with Milli-Q water to remove traces of ethanol, and 1 mL Milli-Q water was added to each tube. The seeds were imbibed at 4 °C for 24 hr to soften the seed coats. Afterwards, the seeds were dissected in ultrapure (Milli-Q) water under a dissecting microscope to obtain the outer seed coats (*i.e.*, the thickened cell wall of oi1). The embryos, endosperm, and inner seed coats (*i.e.*, inner integument layers) were discarded. The outer seed coats were then cut into small pieces (∼1 mm^2^) with razors and transferred into a new 2 mL tube and stored at −80 °C until ss NMR analysis. We used 57 mg of fully hydrated outer seed coat of the wild-type and *tt8* mature seeds.

Ss NMR were performed at 9.4 T (399.86 MHz for ^1^H and 100.55 MHz for ^13^C) on an Agilent VNMRS spectrometer at the Environmental Molecular Sciences Laboratory (EMSL), Pacific Northwest National Laboratory (PNNL), using a homebuilt 4 mm MAS probe tuned to ^1^H/^13^C. The MAS housing (Revolution NMR) was fabricated from Kel-F for minimal ^13^C background signal. The rotors are standard zirconia sleeves with double o-ring Kel-F spacers (Revolution NMR). Carbon chemical shifts were referenced to a secondary standard of the methylene peak of adamantane at 38.48 ppm relative to TMS at 0 ppm. Signal generation of ^13^C was through cross polarization (CP) (Pines *et al*.) with a standard ramped CP pulse sequence using a 3.0 µs ^1^H 90° pulse, a 1 ms contact pulse with a ramped ^1^H RF amplitude, (Metz *et al*., 1994) and a 2 s recycle delay.

SPINAL-64 decoupling (Fung *et al*.) was applied during acquisition at a ^1^H nutation frequency of 83 kHz. The lack of a background signal was verified by performing an experiment under identical conditions on a sample of pure KBr (potassium bromide), the resulting spectrum is shown in **Figure S8**.

### Dry weight measurements of mature embryos and seed coats

Mature pennycress wild-type and *tt8-2bp* seeds, respectively, harvest from six different plants (*i.e.*, six biological replicates) of each genotype were randomly selected. All 12 plants were grown and harvested at the same time, and the seeds were stored in sealed 1.5 mL Eppendorf tubes for about one year before this analysis. From each plant, two sets of 75 seeds were randomly selected from among all the seeds harvested for the measuring whole seed weights and measuring embryo and seed coat weights, respectively. For whole seed weights, the total weights of 75 seeds of each biological replicate were measured on an analytical balance before lyophilized for 24 hr to remove water from the seeds. Immediately after lyophilization, the total weights of 75 seeds were measured again. To obtain embryo and seed coat dry weights separately, the seeds were cleaned and imbibed in the same way as done for the seeds used for the ss NMR analysis. The seeds were dissected in water under a dissecting microscope to separate embryos from seed coats. The embryos and seed coats of each biological replicate were transferred into two different 2 mL Eppendorf tubes. The embryos and seed coats were lyophilized for 24 hr before weighed on an analytical balance. Statistical analyses were done to determine if there is a significant difference in any of the measured dry weights or water contents between wild-type and *tt8-2bp* mature seeds.

### Seed coat permeability measurement

Mature pennycress wild-type and *tt8-2bp* seeds, respectively, harvest from seven different plants (*i.e.*, seven biological replicates) of each genotype were randomly selected. From each plant, three sets of 50 seeds were randomly selected from among all the seeds and transferred into 2 mL Eppendorf tubes for imbibing with Milli-Q water, 20 mg/mL safranin O dissolved in Milli-Q water, and 20 mg/mL toluidine blue O dissolved in Milli-Q water. All the seeds were imbibed in 1 mL of Milli-Q water or dissolved dyes at 4 °C for four days. Afterwards, the dissolved dyes were removed, and the seeds were rinsed with Milli-Q water until the water became clear. Then, ten out of the 50 seeds were randomly sampled from each biological replicate and dissected under a dissecting microscope for observation and counting.

### Seed imbibition rate measurement

Mature pennycress wild-type and *tt8-2bp* seeds, respectively, harvest from seven different plants (*i.e.*, seven biological replicates) of each genotype were randomly selected. From each plant, approximately 200 mg of seeds were weighed on an analytical balance to record the exact weights and transferred into a 2 mL Eppendorf tubes for imbibing in 1 mL Milli-Q water. After water was added to all the tubes, the tubes were vortex briefly and centrifuged at 2,000 *g* for three minutes to submerge all the seeds in water. Then, all the seeds were imbibed at 4 °C for 24 hours. The seeds in each tube were weighed after 1, 2, 3, 4, 6, and 24 hours of imbibition. After each time period, water was removed from all the tubes by pipetting to stop imbibition. Then, the seeds in each tube were patted dry on a piece of KimWipe before weighed on an analytical balance to record the seed weights after imbibition. Immediately after weighing, the seeds were transferred back into the Eppendorf tube and the cap was closed to prevent water loss. After all the seeds were weighed, 1 mL Milli-Q water was added to each tube and the tubes were vortexed and centrifuged to help submerge all the seeds to continue imbibition at 4 °C. The seed imbibition rate (S_i_) after any length of imbibition period described earlier was calculated using the formula S_i_ = (W_i_ – W_b_) / W_b_ where W_i_ and W_b_ represent the seed weights before and after imbibition, respectively. The seed imbibition rates (S_i_) were shown as percentages in **Fig. 9A**. Statistical analyses were done to determine if there is a significant difference in the imbibition rates at any of the timepoint measured between wild-type and *tt8-2bp* mature seeds.

### Seed germination rate measurement after sterilization

The sterilization methods using 70% ethanol and chlorine gas were adapted from published studies with pennycress and Arabidopsis seeds (Ott *et al*., 2021). To sterilize pennycress seeds with 70% ethanol, the seeds were transferred into 2 mL Eppendorf tubes, rinsed twice with 70% ethanol, and then rinsed twice with sterile Milli-Q water. During rinsing, sterile techniques were used, and all the steps were carried out in a biosafety cabinet. For each rinse, 1 mL of 70% ethanol or water was added to each tube and the tube was vortexed for 3 s before the liquid was removed. To sterilize pennycress seeds with chlorine gas, the seeds were transferred into a plastic petri dish or a 24-well plate with lids. Note that the petri dishes or wells cannot be marked with permanent markers because chlorine gas erases the pigments. Sample marking can be done with a tape and pencil. The petri dish or welled plate along with the lid was transferred into a desiccator placed in a fume hood. A 250 mL glass beaker with 150 mL of Clorox® Disinfecting Bleach (7.4% sodium hypochlorite) was also placed in the same desiccator, and a small glass vial with 2 mL of 37% hydrochloric acid was quickly dropped in the glass beaker with forceps. The lid of the desiccator was also quickly closed to prevent the escape of chlorine gas, and the chlorine gas concentration inside the sealed desiccator would reach ∼6.8% (Lindsey Iii *et al*., 2017). After incubating the seeds for desired amount of time, the desiccator was opened to vent for 1 hr. After venting, the lid of the petri dish or welled plate was placed back on. To test seed germination rates after the seeds were sterilized with 70% ethanol or chlorine gas, the seeds were transferred into a well of a sterile 24-well plate immediately after sterilization inside a biosafety cabinet using sterile techniques. The seeds were left inside the biosafety cabinet for 24 hr before 2 mL of sterile Milli-Q water was added to each well.

The plate was wrapped with parafilm, and the seeds were imbibed for 24hr at 4 °C. Finally, the plate was put into the same growth chamber where we growth pennycress plants to start seed germination. To compare germination rates after sterilization by 70% ethanol, 1.5-hr incubation in chlorine gas, and 4-hr incubation in chlorine gas, 25 seeds were randomly selected from five different plants of wild-type and the *tt8* mutant, respectively, and their germination rates were measured after each of the three sterilization methods. We consider the emergence of root radicle as germination, and the number of germinated seeds were counted once a day for eight consecutive days after the germination started. We calculated the seed germination rates of wild-type and *tt8* seeds after different sterilization treatments for statistically analyses.

### Statistical analyses

All student’s t-tests and one-way ANOVA tests were done with R (version 4.3.3) (Team, 2024). The significance threshold is p-value ≤0.05.

For MALDI-MSI analysis, average ion intensities of three biological replicates (i.e., MALDI-MSI data collected from seed sections of three different seeds) were used in comparisons of all the molecular features of interests between wild-type and *tt8-2bp* seed sections at 27 DAP. The log2-transformed average ion intensities of were used in two-tailed student’s t-test to determine if there was a statistically significant difference between wild-type and *tt8-2bp* seeds.

For comparing dry weights of whole seeds, embryos, and seed coats as well as seed water contents, two-tailed student’s t-tests were used for measurements collected from six biological replicates of wild-type and tt8-2bp seeds, respectively.

For comparing seed imbibition rates, seven biological replicates were used for wild type and *tt8-2bp*, respectively. Two-tailed student’s t-tests were done to compare the seed imbibition rates of wild-type and *tt8* seeds after each period of imbibition.

To test how genotype and chlorine gas treatment time affected the germination rates independently and interactively, we performed two-way ANOVA tests using the seed germination rates of wild-type and *tt8-2bp* seeds after 1.5-hour and 4-hour of chlorine gas sterilization on each day after germination started. Five biological replicates were used for wild type and *tt8-2bp*, respectively, for 1.5-hour or 4-hour chlorine gas treatment. The two-way ANOVA tests were done with IBM SPSS Statistics version 30.0 using a mixed effect model where the genotype is the between-subject variable, and the chlorine gas treatment time is the within-subject variable.

To test how 70% ethanol treatment affected germination rates of wild-type vs. *tt8-2bp* seeds, we performed two-tailed student’s t-tests comparing their germination rates on each day after germination started.

To test whether there was significant difference in germination rates of wild-type and tt8-2bp seeds, respectively, on day 8 after different surface sterilization treatments, we performed one-way ANOVA tests. Five biological replicates were used for wild type and *tt8-2bp*, respectively.

## Results

### No anatomical defects were detected in endosperm and embryo development of *tt8-2bp* seeds

In this study, we characterized the anatomy of developing Spring32 wild-type and *tt8-2bp* knockout (KO) mutant seeds at 7-27 DAP. To help understand the results of this study, we will first briefly describe the anatomy of the Spring32 wild-type seed and seed coat. As shown in **Fig. 1A** and **1B**, the embryo of 7 DAP wild-type seeds was at the heart stage and the endosperm was syncytial. Wild-type pennycress seed coats have three ii and oi layers, respectively, where ii1 accumulates PAs (**Fig. 1B** and **1C**). At 27 DAP (**Fig. 1C**), the embryo grew to occupy most of the space inside the seed, and the oi1 cells developed thickened cell wall which eventually became the hardened coat protecting the inner seed compartments from the outer environment.

**Figure 1.**
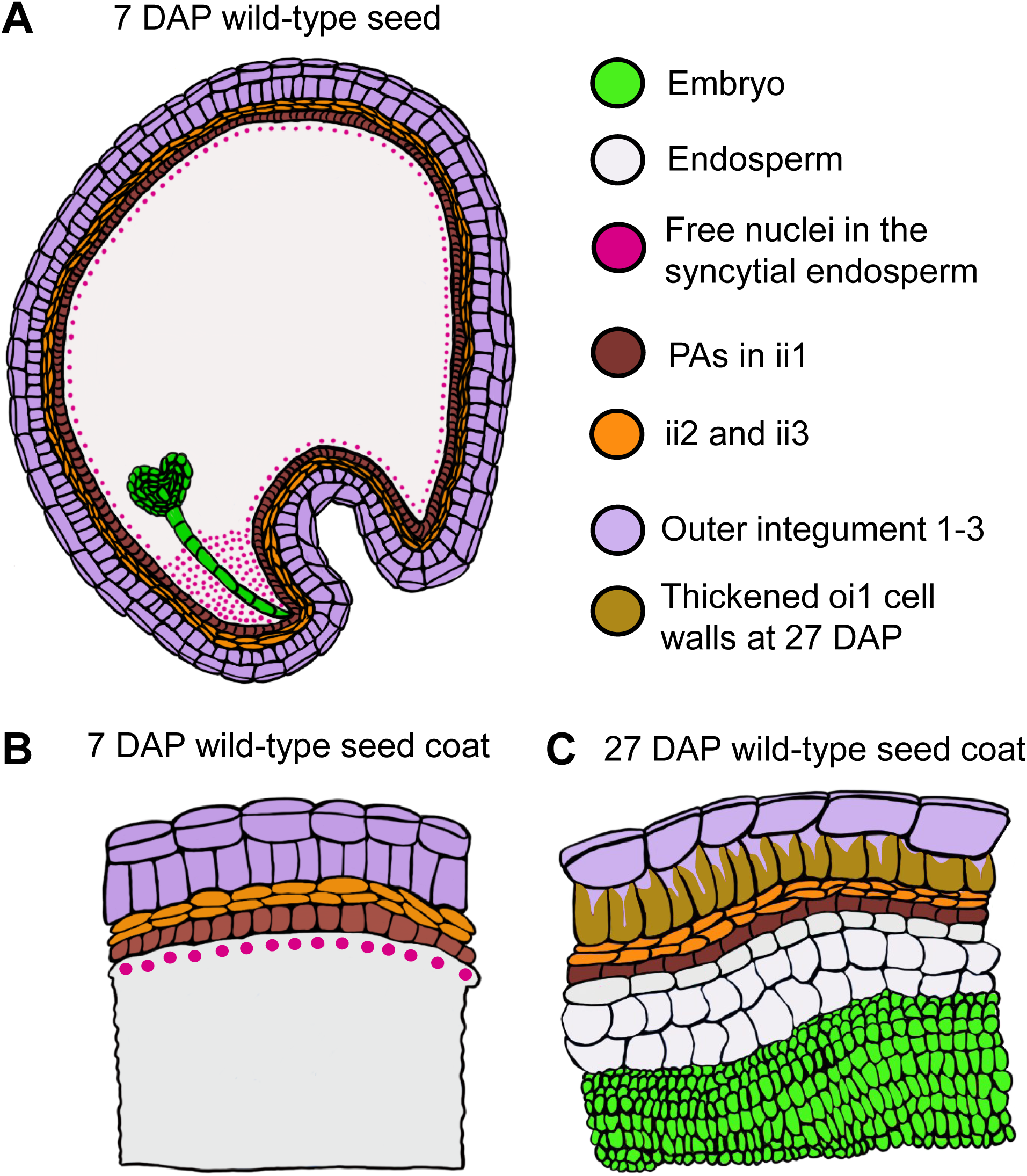
Diagrams depict the anatomy of Spring32 wild-type pennycress seeds, emphasizing seed coat structure at 7 and 27 DAP. (A) A diagram showing anatomy of the cross-section of a 7 DAP pennycress seed. (B, C) Diagrams showing the seed coat cell layers and location of PAs.

Given that *tt8* KO mutations are crucial to pennycress domestication, it is important to assess *tt8* seed traits including any anatomical changes in seed development that could put the crop at risk. Therefore, we examined the seed anatomy of *tt8-2bp* and wild-type seeds at 7, 11, 15, 19, 23, and 27 DAP when pennycress seeds and embryos undergo important transcriptional, metabolic, and morphological changes (Tsogtbaatar *et al*., 2015; Johnston *et al*., 2022). We manually pollinated pennycress flowers of *tt8-2bp* and wild-type plants and counted the day after pollination as 1 DAP (Material and Methods**, Supplementary Fig. S1A**). After harvest, the seeds were immediately fixed using the fixation solution (Material and Methods). Then, we dehydrated the seed samples, embedded them in paraffin, and cut 8-µm sections for histological staining. For analyzing cell structures, we stained the sections with safranin O and alcian blue. Safranin O and alcian blue are cationic dyes mainly used for staining plant cell walls, and can be used together because they have different affinities to various cell wall components (Blokhina *et al*., 2017; Santos *et al*., 2023). Alcian blue primarily stains acidic polysaccharides which are the main components of primary cell walls. In contrast, safranin O has a high affinity to proteoglycans and lignified secondary cell walls (Baldacci-Cresp *et al*., 2020; Pang *et al*., 2023). For each developmental stage, we stained and visualized seed sections from at least three different seeds harvested from three plants.

Under the growth condition of this study (Material and Methods), seed pods and seeds of *tt8-2bp* and wild-type rapidly grew in size until approximately 19 DAP, and the seed pods of both genotypes started to turn yellow at 19 DAP when most leaves were completely senesced (**Supplementary Fig. S1B**, **Supplementary Fig. S2**). At 27 DAP, embryos of wild-type and *tt8-2bp* and seed coats of *tt8-2bp* turned a paler green, suggesting reduced chlorophyll contents (**Supplementary Fig. S2**). Seed coats of 27 DAP wild-type seeds turned reddish-brown, indicating PA oxidation (**Supplementary Fig. S2**).

Based on the stained seed sections at 7, 11, 15, 19, 23, and 27 DAP, the embryo and endosperm of *tt8-2bp* and wild-type seeds appeared to develop at a similar rate and no visible anatomical defects were detected in embryo and endosperm of *tt8-2bp*. The main seed tissues in a stained pennycress seed section were labeled in **Supplementary Fig. S3**. Wild-type and *tt8-2bp* embryos at 7 and 11 DAP were at the heart stage and torpedo stage, respectively (**Fig. 2A**). At 7 DAP, the endosperm of both genotypes was syncytial as the endosperm nuclei were not surrounded by cell wall (**Fig. 2B**). At 11 DAP, the endosperm of both genotypes underwent cellularization and cell wall was visible (**Fig. 2B**). At 15 DAP, wild-type and *tt8-2* embryos grew significantly in size and the cotyledons were completely bent over (**Fig. 2A**, **Supplementary Fig. S2** and **S4**) while the endosperm cells also appeared to have grown bigger (**Fig. 2B**). From 15 to 27 DAP, wild-type and *tt8-2* embryos continued to grow larger and eventually occupied most of the space inside the seed coats (**Fig. 2A**). Meanwhile, the endosperm of both genotypes appeared to gradually decrease in volume and number of cell layers (**Fig. 2A** and **2B**), suggesting that endosperm cells were eliminated to accommodate the growth of the embryo in a way similar to the Arabidopsis endosperm (Huang *et al*., 2023).

**Figure 2.**
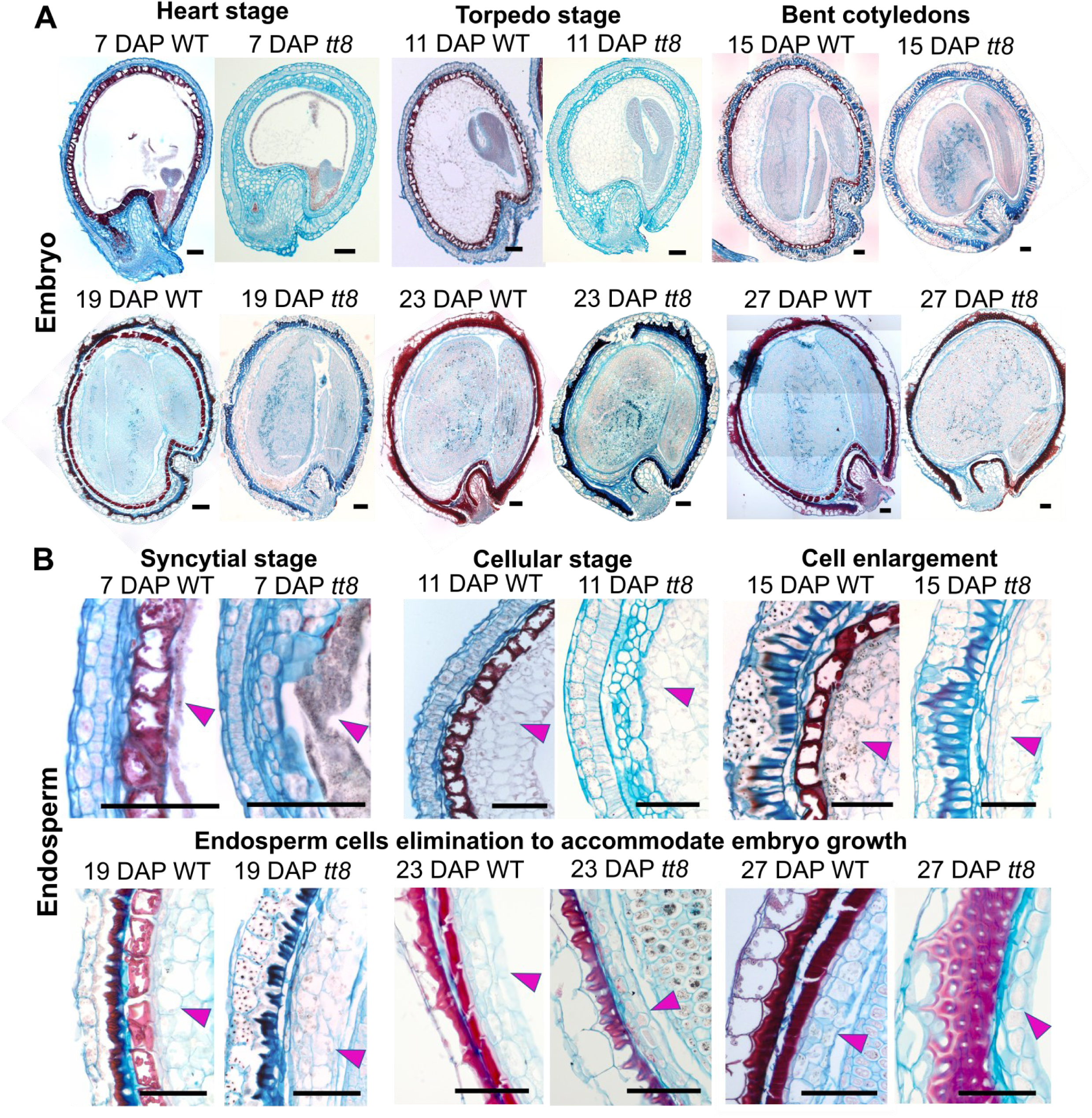
Histological staining of wild-type (WT) and *tt8-2bp* embryo and endosperm shows that the two seed tissues developed at a similar rate at 7-27 DAP. Developing seeds were fixed in 10% formalin and 8-µm paraffin sections were obtained. Rehydrated sections were stained with safranin-O (stains secondary cell wall and nuclei red) and counter-stained with alcian blue (stains acidic polysaccharides of primary cell wall blue). The stained seed sections at 7-27 DAP show that the embryo (A) and endosperm (B) develop at a similar rate. The magenta arrow heads point to the endosperm cells. At least three seeds from three different plants were studies for each developmental stage. Scale bars = 100 µm.

### The ii1 cells of *tt8-2bp* did not accumulate PAs like those of wild-type but stored starch granules

The Arabidopsis seed coat consists of two distinct integuments, *i.e.*, the ii and the oi, comprised of three and two cell layers, respectively (Haughn and Chaudhury, 2005). In Arabidopsis, the innermost layer of ii, ii1, accumulates PAs (Haughn and Chaudhury, 2005) whereas the outmost layer of oi, oi2, accumulates mucilage (Windsor *et al*., 2000). Seed coats of pennycress Spring32 wild-type and *tt8-2bp* seeds contained three layers of ii and oi, respectively (**Fig. 3A**). We labeled the ii and oi cell layers 1-3 based on their distances to the embryo following a similar naming system as in Arabidopsis (**Fig. 3A**).

**Figure 3.**
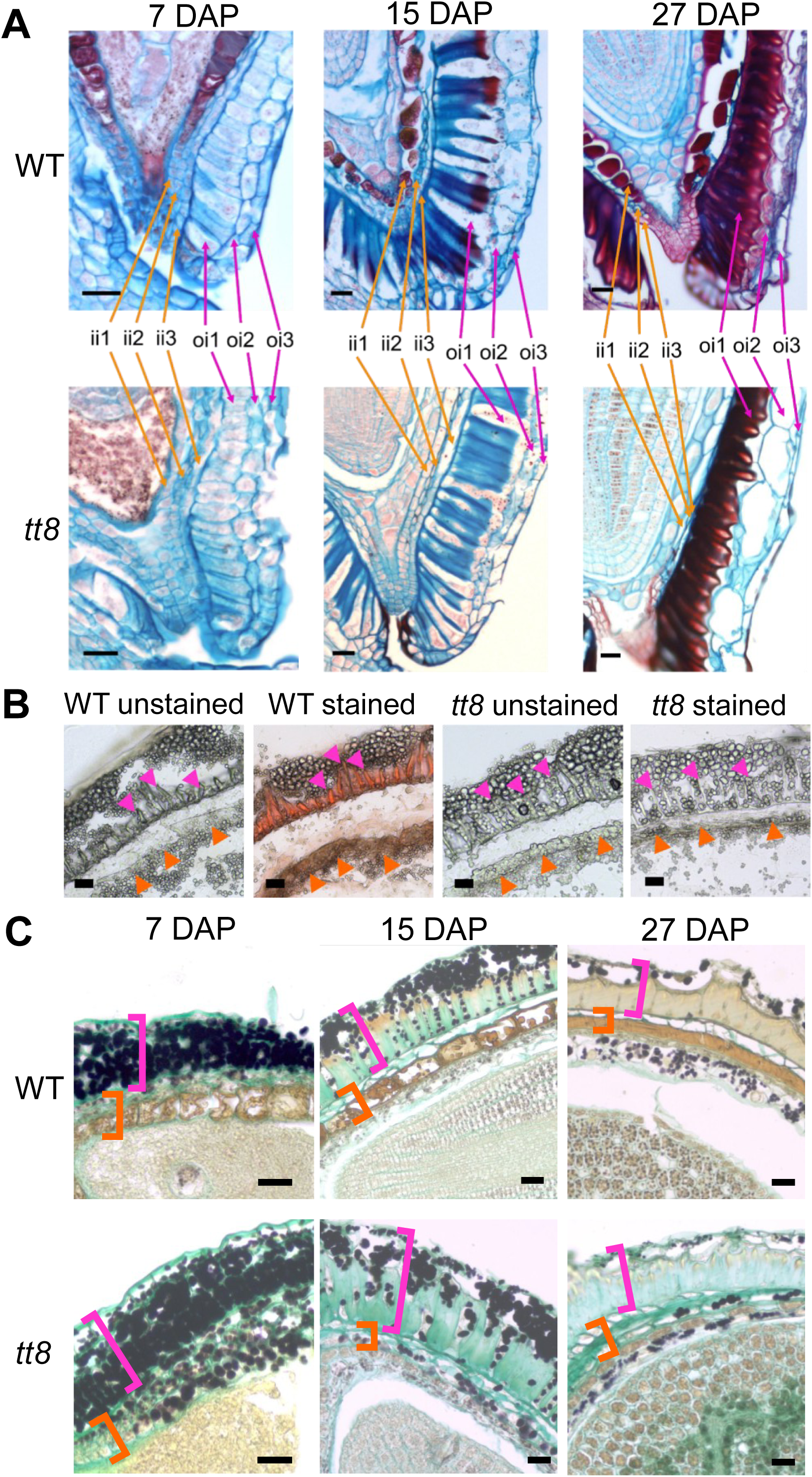
Three cell layers were found in the ii and oi of the Spring32 wild-type (WT) and *tt8-2bp* seed coats where the ii1 cells of WT and *tt8-2bp* accumulate different metabolites during development. (A) Seed sections of WT and *tt8-2bp* at 7, 15, and 27 DAP were stained with safranin-O (stains secondary cell wall and nuclei red) and counter-stained with alcian blue (stains acidic polysaccharides of primary cell wall blue). Note only the WT ii1 cells stained red. (B) Seed cryosections of WT and *tt8-2bp* at 15 DAP were stained with vanillin-HCl which stains PAs reddish-brown. The orange arrowheads point to the ii1 cells and the magenta arrow heads point to the thickened cell wall of oi1 cells. (C) Seed sections of WT and *tt8-2bp* at 7,15, and 27 DAP were stained with alcian blue and lugol’s iodine which stain primary cell wall light turquoise and starch granules black, respectively. The orange brackets mark the ii1-3 and the magenta brackets mark the oi1-3. Note that the ii1 cells of *tt8-2bp* seeds accumulated starch granules. Scale bars = 25 µm.

While wild-type and *tt8-2bp* seeds had the same numbers of integument cell layers, *tt8-2bp* seed coats had drastic changes in PA accumulation in ii1 cells and formation of oi1 cell wall compared to wild type. The pennycress ii and oi cell layers can be distinguished at the micropyle region where the three ii cell layers enclosed endosperm cells adjacent to the embryo root radical whereas oi cell layers left a small opening at the micropyle (**Fig. 3A**).

The wild-type ii1 cells accumulated PAs at 7-27 DAP. When stained with alcian blue and safranin O, the wild-type ii1 cells were filled with metabolites stained dark red which were not observed in the *tt8-2bp* ii1 cells (**Fig. 3A**). Since *TT8* is required for PA synthesis in seed coats, it is likely that the metabolites inside wild-type ii1 cells are PAs. Then, we stained the seed paraffin sections of 15 DAP wild-type and *tt8-2bp* seeds with vanillin-hydrochloric acid (HCl) solution (Materials and Methods). Vanillin-HCl staining is a simple and sensitive method for detecting PAs in plant tissues as vanillin reacts with PAs and forms red compounds (Deshpande *et al*., 1986; Feng *et al*., 2014; Xuan *et al*., 2014). When we compared the paraffin sections of wild-type and *tt8-2bp* seeds at 15 DAP after staining (**Supplementary Fig. S5**), wild-type ii1 cells accumulated yellowish-brown metabolites in both the unstained and vanillin-HCl stained paraffin sections whereas no colored metabolites were detected in the *tt8-2bp* ii1 cells. While no red compounds were observed in vanillin-HCl stained wild-type ii1 cells, we could not rule out that the yellowish-brown metabolites were PAs. It is possible that PAs in the paraffin-embedded wild-type seeds were modified during sample processing and thus cannot react with vanillin-HCl. Therefore, we repeated the vanillin-HCl staining with wild-type and *tt8-2bp* seed cryosections at 15 DAP where no chemicals were used for sample processing (Materials and Methods). After vanillin-HCl staining, red compounds were only observed in ii1 and oi1 cells of wild-type seed cryosections (**Fig. 3B**), suggesting that PAs were accumulated in wild-type but not *tt8-2bp* ii1 and oi1 cells.

The *tt8-2bp* ii1 cells accumulated uncolored granules similar to those accumulated in the oi cells of wild-type and *tt8-2bp* seeds (**Fig. 3B**). Since wild-type and *tt8-2bp* seed coats appeared green during development (**Supplementary Fig. S2**), we hypothesized that their seed coats can perform photosynthesis and store photosynthates in starch granules, *i.e.*, the uncolored granules we observed. Therefore, we stained the paraffin sections of wild-type and *tt8-2bp* seeds with alcian blue and

Lugol’s iodine to visualize primary cell wall and starch granules (Materials and Methods). After staining, the primary cell wall and starch granules were light turquoise and black, respectively (**Fig. 3C**). Starch granules were detected in wild-type and *tt8-2bp* oi cells with the amount of starch granules gradually decreasing from 7 to 27 DAP in both genotypes (**Fig. 3C**). As for the ii cells, starch granules were detected in the ii2 and ii3 layers of wild-type and *tt8-2bp* seeds at 7 DAP but were undetectable at 15-27 DAP (**Fig. 3C**). At 7-27 DAP, wild-type ii1 cells only accumulated PAs whereas *tt8-2bp* ii1 cells accumulated starch granules (**Fig. 3C**).

### Histological analyses of the oi layers indicated PA deficiency in the thickened oi1 cell wall of *tt8-2bp*

In both wild-type and *tt8-2bp* seeds, oi1 cells formed thickened cell wall starting sometime between 11 to 15 DAP (**Fig. 2B**, **Supplementary Fig. S4**). The thickening of oi1 cell wall was uneven as the anticlinal cell walls (*i.e.*, cell walls perpendicular to the seed surface) thickened significantly more than the periclinal cell walls (*i.e.*, cell walls parallel to the seed surface) from 15 to 27 DAP (**Fig. 4A** and **4D**). The oi1 periclinal cell wall adjacent to ii3 had considerably less thickening compared to the anticlinal cell wall whereas the oi1 periclinal cell wall adjacent to oi2 did not show any thickening compared to primary cell walls by 27 DAP (**Fig. 4A** and **4D**). The oi2 and oi3 cell walls in wild-type and *tt8-2bp* seeds did show any thickening at 15-27 DAP (**Fig. 4A**).

**Figure 4.**
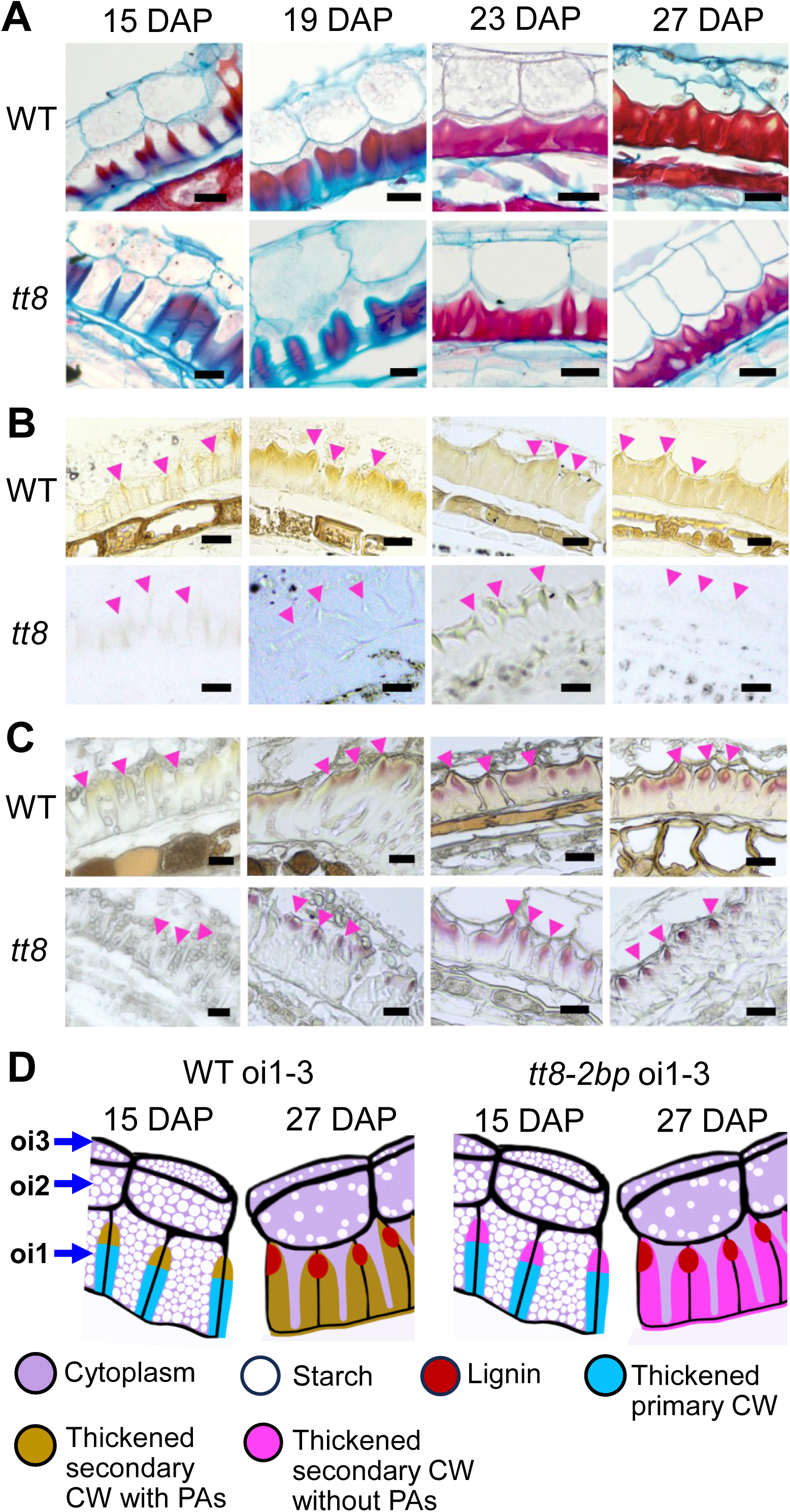
The thickened oi1 cell wall (CW) of Spring32 wild-type (WT) and *tt8-2bp* seeds have different chemical compositions at 15-27 DAP. (A) WT and *tt8-2bp* seed paraffin sections were stained with safranin-O (stains secondary CW and nuclei red) and counter-stained with alcian blue (stains acidic polysaccharides of primary CW blue). (B) WT and *tt8-2bp* seed paraffin sections were imaged without staining. Some brown pigments were observed in the thickened CW of the WT oi1 cells. (C) WT and *tt8-2bp* seed paraffin sections were stained with phloroglucinol-HCl to detect lignin which turned red after staining. (D) Diagrams of oi cell layers depicting anatomical changes in wild-type and *tt8-2bp* oi1 cells from 15 to 27 DAP. The black lines are the thin primary CW. In oi1 cells, the black lines indicate the locations of thin primary CW before CW thickening started. Magenta arrowheads in panel B and C point to the locations of the thickened oi1 CW closest to oi2 cells. Scale bars = 25 μm.

The anatomical changes of oi layers at 15-27 DAP showed some similar patterns in wild-type and *tt8-2bp* seeds. The portion stained red in the thickened oi1 cell walls increased from 15 to 27 DAP, especially in the anticlinal cell walls (**Fig. 4A**). Because alcian blue stains the primary cell wall blue and safranin O stains the secondary cell wall red, this suggests that both wild-type and *tt8-2bp* seeds developed secondary cell wall during the oi1 cell wall thickening from 15 to 27 DAP. We noticed differences in the oi1 thickened cell walls of wild-type vs. *tt8-2bp* when stained with alcian blue and Lugol’s iodine. At 27 DAP, alcian blue stained the *tt8-2bp* thickened oi1 cell wall a light turquoise color (**Fig. 3C**, **Supplementary Fig. S6A**), indicating that acidic polysaccharides such as pectin were present in the secondary cell wall of *tt8-2bp* oi1 cells. This is not surprising since pectin is present in plant primary and secondary cell walls. Interestingly, in 27 DAP wild-type seeds the entire thickened oi1 anticlinal cell wall was not stained by alcian blue but showed a yellowish-brown color similar to PAs accumulated in ii1 cells (**Fig. 3C**, **Supplementary Fig. S6A**). This is consistent with our earlier observation that PAs were accumulated in wild-type ii1 and oi1 cells (**Fig. 3B**), suggesting that the yellowish-brown color of the wild-type oi1 thickened cell wall came from PAs. Therefore, although oi1 cells of wild-type and *tt8-2bp* seeds developed secondary cell walls by 27 DAP, the chemical compositions of their secondary cell walls were different.

We performed additional histochemical analyses to characterize the differences in the thickened oi1 cell wall of wild-type *vs. tt8-2bp* seeds. In unstained seed paraffin sections, the yellowish-brown metabolites (presumably PAs) were again observed in ii1 and the thickened oi1 cell walls of wild-type seeds (**Fig. 4B**). In 15 DAP wild-type seeds, the yellowish-brown metabolites appeared to be most abundant at the end of the oi1 anticlinal cell wall closer to oi2 (**Fig. 4B** and **4D**). As the oi1 anticlinal cell wall gradually thickened, the yellowish-brown metabolites spread more evenly into the oi1 anticlinal cell wall at 23 and 27 DAP (**Fig. 4B** and **4D**). Because lignin is a critical component of secondary cell walls, we stained seed paraffin sections with phloroglucinol-hydrochloric acid (HCl) solution (Materials and Methods), which stains lignin aromatic aldehydes pink or red (Pomar *et al*., 2002). Using phloroglucinol-HCl staining as an indicator, the thickened oi1 anticlinal cell wall started to accumulate lignin between 15 and 19 DAP (**Fig. 4C**). Stained lignin aromatic aldehydes were detected in a small region of the oi1 thickened anticlinal cell walls close to the oi2, and there were no visible differences in lignin accumulation patterns of wild-type *vs*. *tt8-2bp* seeds (**Fig. 4C** and **4D**).

Arabidopsis seed coats accumulate mucilage consisting of complex pectinaceous polysaccharides (Western *et al*., 2000). The seed coat mucilage expands after absorbing water and extrudes to completely cover Arabidopsis seeds (Macquet *et al*., 2007). Ruthenium red has high affinity to pectin, nucleic acids, and calcium-binding proteins (Steeling, 1970; Karpel *et al*., 1981; Cook *et al*., 2013), and is used routinely to detect seed mucilage. We incubated mature seeds of Arabidopsis Col-0, pennycress wild type, and pennycress *tt8-2bp* in ruthenium red solution (Materials and Methods). We only observed mucilage around Arabidopsis Col-0 mature seeds (**Supplementary Fig. S6B**). To verify that there was no mucilage accumulated inside any pennycress seed coat cells, we stained wild-type and *tt8*-2bp seed paraffin sections with ruthenium red (**Supplementary Fig. S6C**). At 7, 15, and 27 DAP, ruthenium red stained the primary cell wall and oi1 secondary cell wall red in wild-type and *tt8-2bp* seeds (**Supplementary Fig. S6C**). We did not find evidence of pennycress seed coat cells filling up with water-soluble, pectin-rich mucilage the same way Arabidopsis seed coat cells do (**Supplementary Fig. S6C**).

Based on the histochemical analyses, we summarized the developmental milestones and important anatomical features of oi cells in pennycress wild-type and *tt8-2bp* seeds (**Fig. 4D)**. By 11 DAP, wild-type and *tt8-2bp* oi cells accumulated abundant starch granules and only developed primary cell wall. Between 15 and 27 DAP, the amount of starch granules gradually decreased in wild-type and *tt8-2bp* oi cells. At 7-27 DAP, oi2 and oi3 cells of wild-type and *tt8-2bp* seeds only had thin primary cell walls. Between 11 and 15 DAP, oi1 cells of wild-type and *tt8-2bp* seeds started to develop thickened secondary cell wall where the anticlinal cell walls thickened more than the periclinal cell walls. The oi1 periclinal cell wall adjacent to the oi2 cells did not show any visible thickening by 27 DAP. In wild-type seeds, the thickened oi1 cell walls gradually accumulated PAs and potentially other flavonoids from 15 to 27 DAP. In *tt8-2bp* seeds, oi1 cells developed thickened secondary cell wall without PAs at 15-27 DAP. Between 15 and 19 DAP, the thickened oi1 anticlinal cell walls of wild-type and *tt8-2bp* started to accumulate lignin in small areas close to oi2 and the areas of lignin accumulation did not expand by 27 DAP. Since the cell wall of oi2 and oi3 did not show any thickening by 27 DAP, it is most likely that the thickened cell wall of oi1 eventually became the hardened protective coat of mature pennycress seeds.

### *tt8-2bp* seeds had decreased PAs in the seed coat and increased small sugars and fatty acids in the whole seed

We performed spatial metabolomic analysis on 27 DAP wild-type and *tt8-2bp* seed cryosections using matrix-assisted laser desorption ionization-mass spectrometry imaging (MALDI-MSI) to corroborate the histological staining results and explore additional changes in chemical compositions of *tt8-2bp vs.* wild-type seeds. We obtained mass spectrometry images of seed cryosections at a high spatial resolution of 25 x 25 μm^2^ per pixel (Materials and Methods). After overlaying the mass spectrometry image with the corresponding brightfield micrograph, we ascribed ion signals of different molecular features to specific seed compartments. The molecular features detected by MALDI-MSI were identified by measuring mass-to-charge ratios (m/z) of intact molecules and thus cannot differentiate molecules with the exact same mass (mass isomers), *i.e.*, when molecules with the same molecular formula are ionized simultaneously. For example, the molecular feature C_6_H_12_O_9_S can represent D-Galactose 6-sulfate, D-Glucose 6-sulfate, and/or 6-Deoxy-6-sulfo-D-gluconate with m/z of 259.0129. Thus, the relative abundance of a molecular feature in a seed section represents the average abundance of all the ionized molecules with the same m/z, which was measured by the ion signal intensity in MALDI-MSI.

We searched for PAs, PA precursors, precursors of main cell wall components such as cellulose, pectin, and lignin, and main components of seed storage nutrients such as polysaccharides, fatty acids, and amino acids in the MALDI-MSI datasets targeting small molecules and lipids generated under positive and negative ion modes (Materials and Methods). All the molecular features of interest were found in the small molecule and lipid datasets generated under the negative ion mode. For quantitative comparisons, we used the average ion signal intensities calculated based on the signal localizations in wild-type and *tt8-2bp* seed sections, respectively (Materials and Methods). A brightfield micrograph with labeled seed compartments is shown in **Supplementary Fig. S7U**. There were no significant differences between the total pixel numbers of any seed compartment when comparing wild-type vs. *tt8-2bp* seed sections (**Supplementary Fig. S7V**).

Ion signals of proanthocyanidin A2 (C_30_H_24_O_12_) and several other aromatic compounds (C_15_H_12_O_6_) were found exclusively in wild-type seed coats whereas digalacturonate and glycyrrhizin (C_12_H_18_O_13_) and some sugar acids and sugar acid derivatives (C_6_H_10_O_7_) were found exclusively in seed coats of wild type and *tt8-2bp*. Proanthocyanidin A2 ion signal was only detectable in the ii layers and thickened oi1 cell wall of wild-type seed coats (**Fig. 5A-C**), supporting histological staining results of PAs in the ii1 cells and thickened oi1 cell wall of wild-type seeds. The molecular feature C_15_H_12_O_6_ had 14 candidate aromatic compounds including dihydrokaempferol, a flavonoid upstream of PA biosynthesis, and fustin, a bioactive flavonoid, (Roy *et al*., 2022), and their ion signals were also only detectable in wild-type samples (**Supplementary Fig. S7B**). This suggests that the *tt8-2bp* mutation potentially also affected the production of other aromatic compounds in seed coats. Digalacturonate and glycyrrhizin were the only two candidate molecules identified for C_12_H_18_O_13_. Digalacturonate is a pectin degradation product whereas glycyrrhizin is a triterpene compound stored in vacuoles (Kato *et al*., 2022). Based on the seed coat cell wall localization of C_12_H_18_O_13_ ion signals in wild-type and *tt8-2bp* samples (**Supplementary Fig. S7F**), the molecule was mostly likely digalacturonate. The sugar acids and sugar acid derivatives (C_6_H_10_O_7_) have 20 candidate molecules of which many were pectin precursors or pectin degradation products such as D-galacturonate, D-tagaturonate, D-fructuronate, *etc*. The ion signals of C_6_H_10_O_7_ have the same localizations as C_12_H_18_O_13_ (**Supplementary Fig. S7H**), suggesting that the ion signals of C_6_H_10_O_7_ potentially also came from pectin degradation. The average ion intensities of C_6_H_10_O_7_ and C_12_H_18_O_13_ showed no significant difference between wild-type and *tt8-2bp* samples (**Supplementary Fig. S7G** and **S7I**). Since all the seeds used for the MALDI-MSI were grown and harvested at the same time and were all storage at −80 °C before analysis, the rate of pectin degradation should be similar in wild-type and *tt8-2bp* seeds. The lack of significant differences in C_6_H_10_O_7_ and C_12_H_18_O_13_ in wild-type *vs*. *tt8-2bp* samples suggests that the total pectin contents of the wild-type and *tt8-2bp* seed coats may not be significantly different, either. The monomer of cellulose is β-D-glucose (C_6_H_12_O_6_), and in the MALDI-MSI dataset C_6_H_12_O_6_ either had low ionization efficiency or too much background signal under the positive and negative ion modes, respectively. All the molecular features representing lignin precursors or monomers showed completely different ion signal patterns under positive *vs.* negative ion modes, so we cannot extract meaningful information from these data.

**Figure 5.**
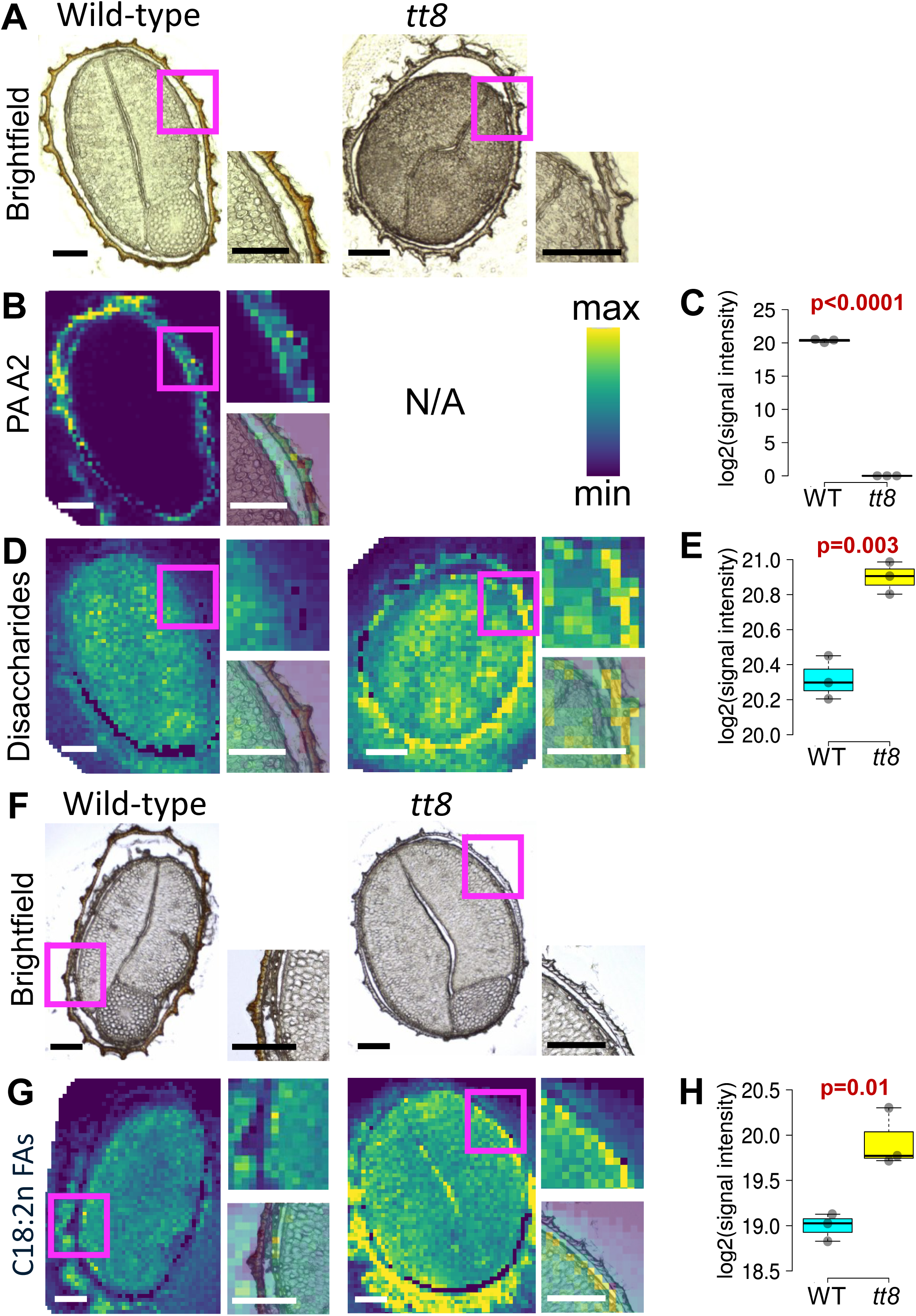
Spatial metabolomic analysis (MALDI-MSI) of small molecules and lipids in Spring32 wild-type (WT) and *tt8-2bp* seed cryosections at 27 DAP. (A, F) Representative images of brightfield micrographs of seed cryosections used for MALDI-MSI for the detection of small molecules and lipids, respectively, under the negative ion mode. A blowup image of the indicated seed region (magenta box) is provided to show the spatial separation of seed coat and embryo. (B, D, G) Heatmaps showing the ion signal intensities of proanthocyanidin A2 (PA A2, C_30_H_24_O_12_), disaccharides (C_12_H_22_O_11_), and C18:2n octadecadienoic acid (C18:2n FAs, C_18_H_32_O_2_) detected in the same seed sections show in panel A and F. The heatmaps of the three different molecular features are displayed in the same color scale with deep blue representing the min value and yellow representing the max value without any data transformation. The min and max values are the same in the heatmaps of WT and *tt8-2bp* seed sections of the same molecular feature. A blowup image of the indicated seed region (magenta box) is shown to the upper right of each heatmap, and an overlaid image of the blowup heatmap adjusted to 50% transparency and the brightfield micrograph is shown to the lower right of each map. Scale bars = 200 µm. Proanthocyanidin A2 is the only molecule identified for C_30_H_24_O_12_ with m/z of 575.1195. There are 36 different disaccharides such as sucrose, lactose, maltose, etc., identified for C_12_H_22_O_11_ with m/z of 341.1089. Three C18:2n octadecadienoic acids including linoleic acid, 18:2(9Z,11E), and (6Z,9Z)- octadecadienoic acid were identified for C_18_H_32_O_2_ with m/z of 279.2330. (C, E, H) Boxplots of log2-transformed average ion signal intensities of each molecular feature detected in three wild-type and *tt8* seed sections, respectively. The p-values calculated by two-tailed student’s t-tests for comparing the log2-transformed average ion signal intensities between WT and *tt8-2bp* seed sections are shown above the boxplots. The gray dots represent values of different seed sections.

Additionally, we searched for metabolites representative of seed storage compounds including polysaccharides, major fatty acids of storage lipids, and amino acids and their derivatives. The MALDI-MSI performed here was customized to identify small molecules and lipids, so proteins were not analyzed. Ion signals of many disaccharides (C_12_H_22_O_11_) and trisaccharides (C_18_H_32_O_16_) were localized to embryos and seed coats of wild-type and *tt8-2bp* samples (**Fig. 5D**, **Supplementary Fig. S7D**). The average ion intensities of disaccharides and trisaccharides were significantly increased in *tt8-2bp* samples by 1.5-fold and 1.6-fold, respectively (**Fig. 5E**, **Supplementary Fig. S7E**). There were 36 and 14 candidate molecules identified for C_12_H_22_O_11_ and C_18_H_32_O_16_, respectively. Disaccharides such as sucrose and maltose are important starch precursor and degradation product, respectively (Hill *et al*., 2003; Stettler *et al*., 2009; Andriotis *et al*., 2010), and trisaccharides such as raffinose and maltotriose are known to accumulate during seed desiccation and starch degradation, respectively (Li *et al*., 2017a; Li *et al*., 2017b). As for fatty acids, we found molecular features representing the most abundant fatty acids in seeds including linoleic acid (C_18_H_32_O_2_), oleic acid (C_18_H_34_O_2_), palmitic acid (C_16_H_32_O_2_), and α-Linolenic acid (C_18_H_30_O_2_) (**Fig. 5F** and **5G**, **Supplementary Fig. S7N-T**). Except for palmitic acid, molecular features representing the other fatty acids had additional candidate molecules which are fatty acids with the same number of carbons and double bonds. Among all the molecular features, C_18_H_34_O_2_ representing oleic acid and other C18:1n fatty acids had a moderate increase in average ion signal intensities in *tt8-2bp vs*. wild-type samples with p-value=0.056 and a 1.3-fold increase (**Supplementary Fig. S7P**). The molecular features representing linoleic acid (C_18_H_32_O_2_, C18:2n fatty acids), palmitic acid (C_16_H_32_O_2_), and α-Linolenic acid (C_18_H_30_O_2_, C18:3n fatty acids) had significantly increased average ion intensities in *tt8-2bp vs*. wild-type samples with p-values≤0.05 and fold changes of 1.9-2.0 (**Fig. 5H**, **Supplementary Fig. S7R** and **S7T**). Additionally, we find two amino acid derivatives, sinigrin and glutathione, with 1.8-fold increase of average ion signal intensities in *tt8-2bp* seed sections compared to those of wild-type (**Supplementary Fig. S7K** and **S7M**). Sinigrin and glutathione are the only metabolites identified for C_10_H_17_NO_9_S_2_ and C_10_H_17_N_3_O_6_S, respectively, and their ion signals were mainly located in the embryos (**Supplementary Fig. S7J** and **S7L**). Sinigrin and glutathione are bioactive molecules in plant seeds with known antimicrobial (Rangkadilok *et al*., 2002; Bischoff, 2021) and antioxidant function (Koramutla *et al*., 2021), respectively, and thus can be important for seed vigor and germination in the field. Taken together, the MALDI-MSI results showed that *tt8-2bp* knockout mutation likely affected not only the seed coat development and chemical compositions but also the accumulation of major seed storage compounds including polysaccharides and fatty acids in embryos. The increase of sinigrin and glutathione in *tt8-2bp* seeds suggest that *tt8* KO mutant seeds may have different responses to biotic and abiotic stresses compared to wild-type seeds in the field.

### Outer seed coats of mature *tt8-2bp* seeds had reduced aromatic compounds and cell wall polysaccharides

Based on the histological staining and MALDI-MSI analyses of 27 DAP seeds, there appeared to be a significant reduction in PAs in both ii1 and thickened oi1 cell wall of *tt8-2bp* seeds without significant impacts on cell wall pectin or lignin contents. The lignified secondary cell wall of oi1 forms the protective outer seed coat enclosing endosperm and embryo in wild-type and *tt8-2*bp mature seeds and can thus affect seed quality and germination. Therefore, we characterized the outer seed chemical compositions of wild-type and *tt8-2bp* mature seeds by ss NMR analysis (Materials and Methods). Ss NMR is complementary to MALDI-MSI as MALDI-MSI only measured small molecules and lipids whereas ss NMR is better at measuring components of large polymers such as lignin, PAs, cellulose, and pectin. We used 57 mg fully hydrated outer seed coats of wild type and *tt8-2bp* for ss NMR. We were not able to measure the water contents of fully hydrated out seed coats due to technical challenges. For quantitative comparisons, we scaled the ss NMR spectra of wild-type and *tt8-2bp* to the peak at 176 ppm representing the carbonyl groups in D-galacturonate and homogalacturanan (**Tab. 1**, **Fig. 6A**). D-galacturonate and homogalacturanan are main components of pectin and were selected for spectrum normalization because pectin was detected in the thickened oi1 cell wall (**Supplementary Fig. S6C**) and were likely of similar abundance in oi layers of wild type and tt8-*2bp* based on MALDI-MSI results (**Supplementary Fig. S7F-I**).

**Figure 6:**
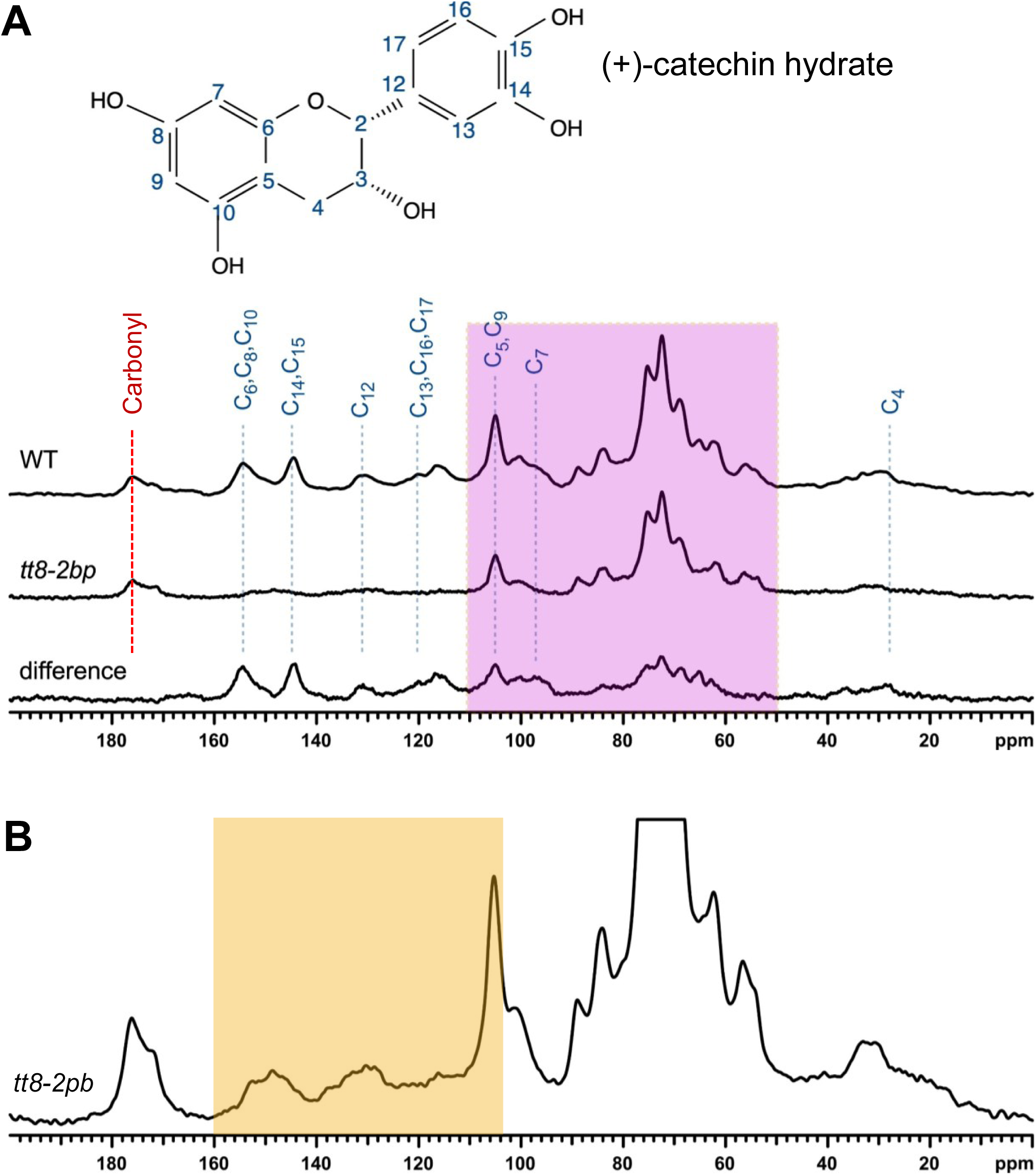
1^3^C CP-MAS spectra of wild-type (WT) and *tt8-2bp* mature outer seed coat show drastic reduction in aromatic compounds in *tt8-2bp* seed coats. (A) Spectra of WT and *tt8-2bp* after scaling peak intensities to the 176-ppm peak, representing the carbonyl group of D-galacturonate and homogalacturanan, main components of pectin. The difference spectrum was obtained by subtracting the peak intensities of *tt8-2bp* spectrum from WT spectrum. (+)-catechin hydrate was identified as the main aromatic compound contributing to the differences between WT and *tt8-2bp* at 50-110 ppm. The molecular structure was shown above the spectra where the numbers correspond to the C number labeled in the spectra below. The pink box highlights peak groups at 50-110 ppm representing chemical groups of cellulose, hemicellulose, and other general carbohydrates. (B) Zoomed-in *tt8-2bp* spectrum emphasizes peak groups at 104-160 ppm (orange box) where chemical groups of aromatic compounds including lignin and PAs are located.

**Table 1.** Assignment of peaks in ^13^C CP/MAS NMR spectra of pennycress outer seed coat.

| Chemical shift (ppm) | Assignment |
| --- | --- |
| 176 | Carbonyl of D-galacturonate and homogalacturanan |
| 154 | C<sub>6</sub>, C<sub>8</sub>, C<sub>10</sub> |
| 144 | C<sub>14</sub>, C<sub>15</sub> |
| 130 | C<sub>12</sub> |
| 120 | C<sub>13</sub>, C<sub>16</sub>, C<sub>17</sub> |
| 105 | C<sub>5</sub>, C<sub>9</sub> |
| 96 | C<sub>7</sub> |
| 98-158 | Aromatic compounds |
| 50-110 | Cellulose, hemicellulose, and general carbohydrates C<sub>2</sub>, C<sub>3</sub>, C<sub>5</sub> |
| 29 | C<sub>4</sub> |

The spectra of wild-type and *tt8-2bp* showed no prominent signal from very-long-chain fatty acids, the main component of cuticle and wax, which are represented by a sharp peak at ∼30 ppm. This suggests that very-long-chain fatty acids are not major components of mature pennycress outer seed coats. To identify the main differences in chemical compositions of wild-type *vs*. *tt8-2bp* outer seed coat, a difference spectrum was calculated (**Fig. 6A**). The main differences were found among peaks representing aromatic compounds (98-158ppm) and polysaccharides (50-110 ppm) where the wild-type spectrum had higher peak intensities than the *tt8-2bp* spectrum. The annotated peaks and peak regions are listed in **Tab. 1**. After aligning the wild-type and difference spectra of aromatic compounds, a close match was found with the known NMR spectrum of (+)-catechin hydrate (Martínez-Richa and Joseph-Nathan, 2003). (+)- catechin hydrate is a hydrated form of catechin, a major PA monomer (Yu *et al*., 2022). We noted that the C5,C9 and C7 peaks of (+)-catechin hydrate overlapped with the range of peak groups representing polysaccharides (**Fig. 6A**), suggesting that the peak intensities of C5,C9 and C7 peaks likely reflected chemical groups in (+)-catechin hydrate and other polysaccharides. In *tt8-2bp* spectrum, the peak intensities of aromatic compounds were very low compared to wild-type spectrum (**Fig. 6A**), but they are detectable after zoomed (**Fig. 6B**). One prominent peak is the one coinciding with the C5,C9 peak of (+)-catechin hydrate, indicating that part of the signal came from polysaccharides in *tt8-2bp* outer seed coats. Lignin was detected in small areas of thickened oi1 cell walls close to the oi2 in 27 DAP wild-type and *tt8-2bp* seeds (**Fig. 4C**) whereas PAs were undetectable in 27 DAP *tt8-2bp* seed coats (**Fig. 6B**), suggesting lignin to be the main aromatic compound in *tt8-2bp* seed coats. Therefore, in *tt8-2bp* spectrum the low intensity peaks representing aromatic compounds excluding the ones overlapped with polysaccharides likely came from lignin (**Fig. 6B**). As for polysaccharides (50-110 ppm), wild-type spectrum had higher peak intensities than *tt8-2bp* spectrum (**Fig. 6A**), suggesting that wild-type outer seed coats had more polysaccharide cell wall components such as cellulose, hemicellulose, *etc*. In summary, ss NMR analysis showed that the main aromatic compounds in mature outer seed coats of wild type and *tt8-2bp* were PAs and lignin, respectively. The fully hydrated outer seed coat of *tt8-2bp* had reduced PAs and cell wall polysaccharides compared to wild type.

### Mature *tt8-2bp* seeds had decreased seed coat dry weights and increased embryo dry weights

Histological staining, MALDI-MSI, and ss NMR analyses showed that developing and mature *tt8-2bp* seed coats had reduced PAs and cell wall polysaccharides compared to wild-type seed coats. Thus, we tested whether mature *tt8-2bp* seeds lost water faster than wild-type seeds as PA deficiency may allow water inside the seeds to escape faster. We measured weights of mature wild-type and *tt8-2bp* seeds from six plants (*i.e.*, six biological replicates) per genotype before and after dehydration (Materials and Methods). The seed water contents were calculated as percentages of seed weights before dehydration. There was no significant difference in seed weights before or after dehydration nor in water contents when comparing wild-type and *tt8-2bp* seeds (**Fig. 7A** and **7B**). However, there may still be differences in embryo or seed coat dry weights of wild-type *vs*. *tt8-2bp* seeds. We then measured the dry weights of embryos and seed coats of wild-type and *tt8-2bp* seeds using six biological replicates, respectively. Mature *tt8-2bp* seeds had significantly lower seed coat dry weights yet significantly higher embryo dry weights compared to wild-type seeds (**Fig. 7C**). This suggests that mature *tt8-2bp* seeds had altered total nutrient partitioning between embryos and seed coats compared to wild-type seeds. To quantify the changes in total nutrient partitioning, we calculated the embryo-to-seed coat ratios by dry weights for wild-type and *tt8-2bp* seeds. Mature *tt8-2bp* seeds had significantly higher embryo-to-seed coat ratios than wild-type seeds (**Fig. 7D**). Since no significant difference was found in whole seed dry weights, we can assume that the total nutrients measured by dry weights fixed in mature seeds including storage compounds (*e.g.*, starch, lipids, and proteins) and structural components (*e.g.*, cell wall fibers) were about equal for wild type and *tt8-2bp*. The embryo-to-seed coat ratios were 2:1 and 3:1 for wild-type and *tt8-2bp* seeds, respectively, indicating that on average 66% and 75% of total nutrients in wild-type and *tt8-2bp* seeds, respectively, were fixed in embryos. Therefore, there was on average a 9% increase in the total nutrients fixed in *tt8-2bp* mature embryos compared to those of wild type.

**Figure 7:**
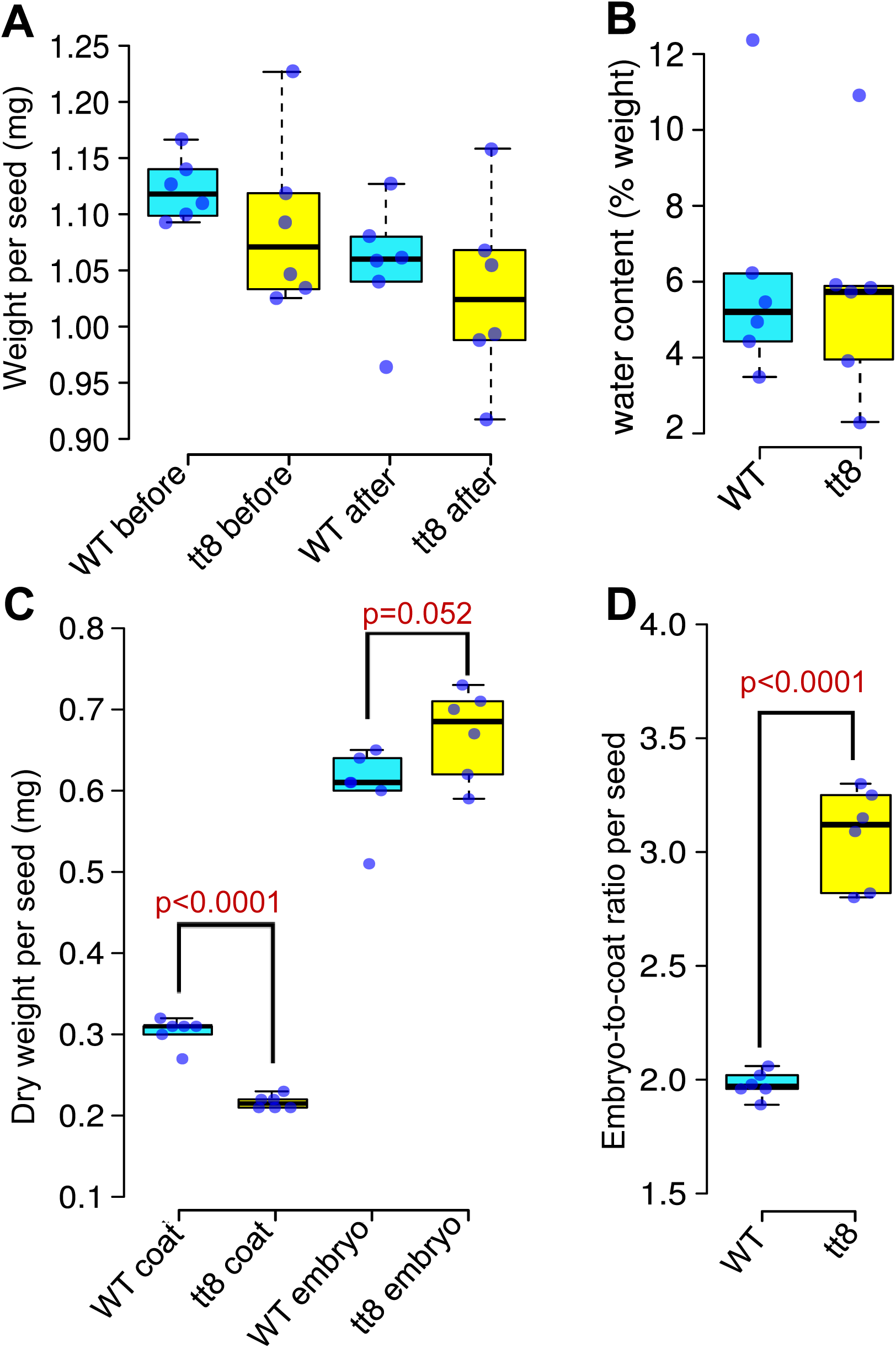
Mature *tt8-2bp* seeds have significantly reduced seed coat dry weights and increased embryo dry weights and embryo-to-seed coat ratios compared to wild-type (WT) seeds. All measurements were done for six biological replicates of 75 seeds per biological replicate. (A) Average weight per seed of WT and *tt8-2bp* before and after dehydration. No statistically significant difference in seed weights of WT *vs. tt8* seeds was found. (B) Average water content per seed as percentage seed weights before dehydration. No statistically significant difference in water contents was found in WT *vs. tt8-2bp* seeds. (C) Average dry weights of seed coat and embryo per seed after dehydration. (D) Ratios of embryo-to-seed coat dry weights calculated based on values in panel-C. P-values are calculated by two-tailed student’s t-test with a threshold of p≤0.05.

### Mature *tt8-2bp* seeds had increased seed coat permeability toward water-soluble dyes

With decreased PAs, cell wall polysaccharides, and dry weights, *tt8-2bp* seed coats may have increased seed coat permeability towards water soluble molecules compared to wild-type seed coats. To assess the seed coat permeability, we soaked wild-type and *tt8-2bp* seeds from seven different plants *(i.e.*, seven biological replicates) per genotype in 20 mg/mL safranin O and toluidine blue O, respectively (Materials and Methods). After incubation at 4 °C for four days, there were significantly higher percentages of *tt8-2bp* seeds with embryos stained by safranin O and toluidine blue O (**Fig. 8A**). The *tt8-2bp* embryo regions near the micropyle and chalazal were most heavily stained (**Fig. 8B**), indicating that dyes permeated through *tt8-2bp* seed coats at these two regions. To further investigate the increased *tt8-2bp* seed coat permeability, we compared the seed coat anatomy of 27 DAP wild-type and *tt8-2bp* seeds at the micropyle and chalazal regions (**Fig. 8C**). The micropyle and chalazal of wild-type and *tt8-2bp* seeds were not completely enclosed with the thickened oi1 cell wall, exposing the ii layers to the outer environment (**Fig. 8D**). The PA-accumulating ii1 cells of wild-type seeds wrapped around the endosperm cells at the micropyle and chalazal regions, forming a barrier protecting the endosperm and embryo (**Fig. 8D**). In contrast, the ii1 cells of *tt8-2bp* seeds only had primary cell wall except for a small number of cells at the micropyle where their cell walls stained red (**Fig. 8D**). It is likely that the thin-walled ii cells at the micropyle and chalazal regions of *tt8-2bp* seeds cannot effectively block water soluble molecules from passing through their primary cell walls, contributing to the increased seed coat permeability.

**Figure 8.**
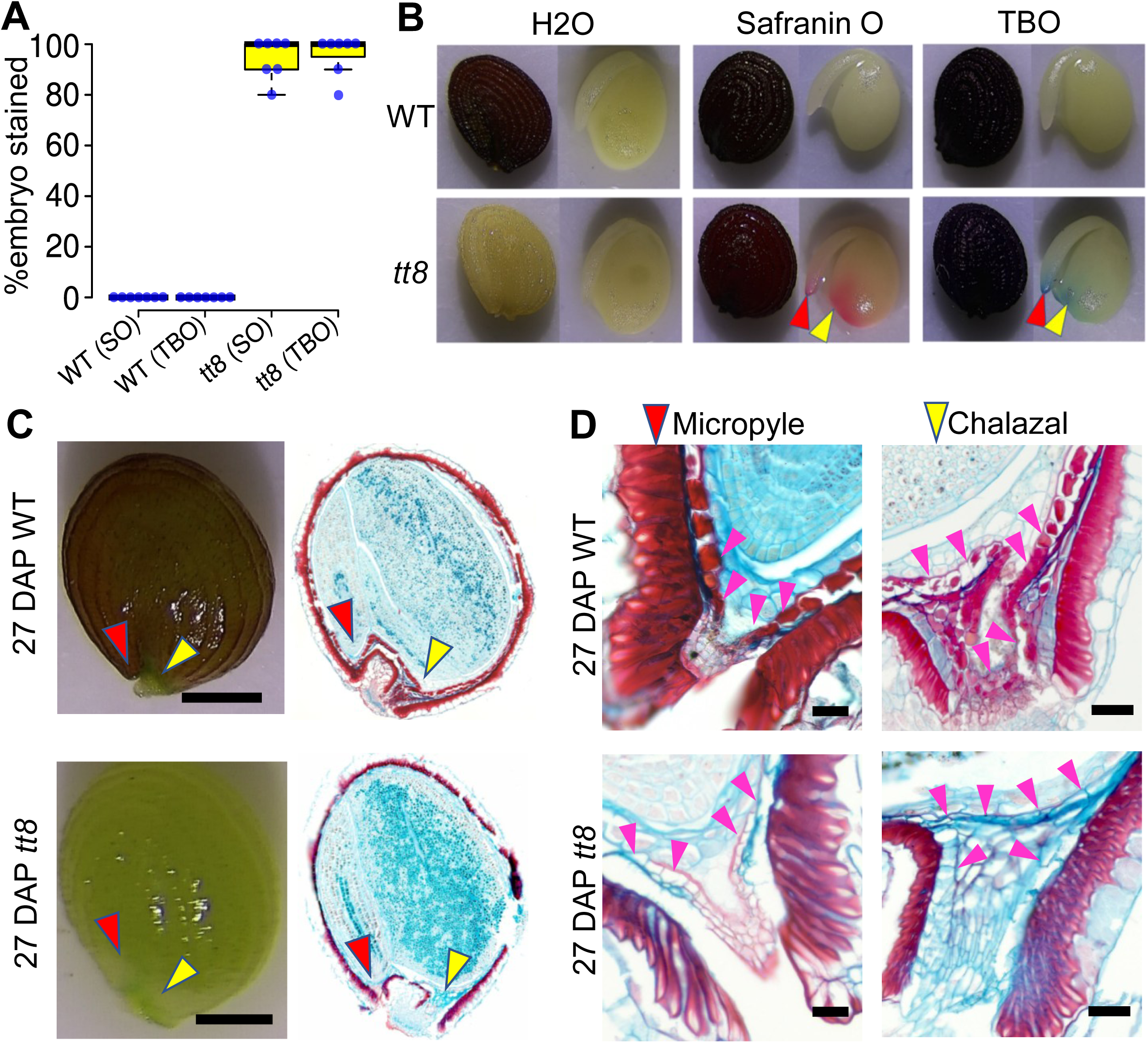
Assessing the seed coat permeability of Spring32 wild-type (WT) and *tt8-2bp* mature seeds. (A) Mature WT and *tt8-2bp* seeds, respectively, were submerged in water, safranin O (SO) (20mg/mL), or toluidine blue O (TBO, 20mg/mL) for four days at 4 °C in dark. *tt8-2bp* seeds were most heavily stained at the micropyle (red arrowhead) and chalaza (yellow arrowhead). (B) Quantification of stained *tt8-2bp* and WT seeds from six biological replicates were used (*i.e.*, seeds from six different plants of each genotype). (C) Images of intact WT and *tt8-2bp* seeds at 27 DAP and micrographs of the seed cross-sections at 27 DAP stained with alcian blue and safranin O. Red and yellow arrowheads point to micropyle and chalaza regions. Scale bars = 500 µm. (D) WT and *tt8-2bp* seed sections of seeds at 27 DAP stained with alcian blue and safranin O demonstrate that the PA-accumulating ii1 cells in WT at the micropyle and chalaza regions (magenta arrowheads) potentially contributed to the lower permeability of WT seed coats. Scale bars = 50 µm.

### Mature *tt8-2bp* seeds had increased imbibition rates and unchanged seed coat gas permeability toward chlorine

Due to altered seed coat chemical compositions and permeability of mature *tt8-2bp* seeds, there were potential changes in the seed imbibition rates and gas permeability. To measure seed imbibition rates, we submerged seeds from seven different plants (*i.e.*, seven biological replicates) of wild type and *tt8-2bp*, respectively, for 1, 2, 3, 4, 6, and 24 hours. The seed imbibition rates at each time point were calculated as percentages of gained water weights over the seed weights before imbibition (Materials and Methods). At 1, 2, 3, and 4 hours after imbibition, *tt8-2bp* seeds had significantly higher imbibition rates than wild-type seeds (**Fig. 9A**). However, at 6 and 24 hours of imbibition, there was no significant difference in imbibition rates between wild-type and *tt8-2bp* seeds (**Fig. 9A**). This suggests that *tt8-2bp* seeds absorbed water faster in the early phase of imbibition but there was not significant difference in the total amount of water absorbed by seeds of the two genotypes. After 24 hours of imbibition, *tt8-2bp* and wild-type seeds on average absorbed water that were 47.3% and 47.9% of their weights before imbibition, respectively.

**Figure 9.**
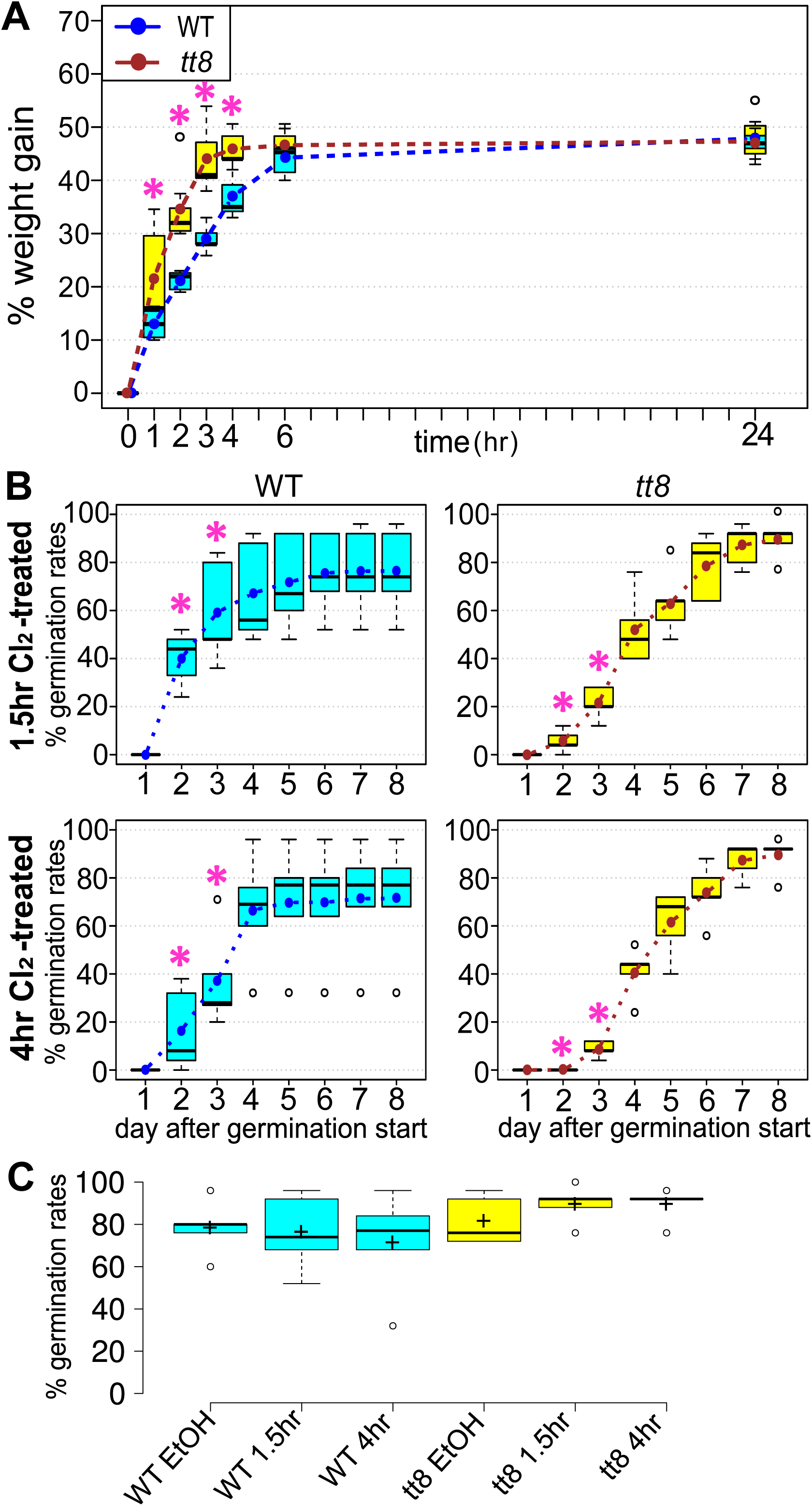
Boxplots of Spring32 wild-type (WT) and *tt8-2bp* (*tt8*) seed imbibition rates and germination rates after sterilization. (A) Boxplots of seed imbibition rates measured by % weight gain after submerging seeds in water. Asterisks indicate statistically significant difference (p-value ≤0.05 between the imbibition rates of WT and *tt8* based on two-tailed student’s t-tests. (B) Seed germination rates after sterilizing seeds with 1.5-hour and 4-hour chlorine gas treatment, respectively. Asterisks indicate statistically significant difference (p-value ≤0.05 between the germination rates of WT and *tt8* based on two-way ANOVA tests. Dotted lines in A-C connect mean germination rates of each day. (C) Seed germination rates on day 8 after germination started of WT and *tt8* seeds treated by 70% ethanol (EtOH), 1.5-hour chlorine gas (1.5hr), and 4-hour chlorine gas (4hr), respectively. No statistically significant difference was found between the seed germination rates based on one-way ANOVA tests with significance threshold of p-value ≤0.05. The mean germination rate was marked by “+”.

To test gas permeability, we performed chlorine gas sterilization on mature wild-type and *tt8-2bp* seeds and compared their germination rates afterward. Chlorine gas sterilization has been standardized in Arabidopsis and can reduce seed germination rates when applied for too long or at too high concentrations of chlorine gas (Lindsey Iii *et al*., 2017), and it was recommended to sterilize Arabidopsis seeds for 1 hour at 6.1% chlorine gas concentration. Surface sterilization with 70% ethanol did not affect seed germination rates of wild-type and *tt2* KO mutant seeds (Ott *et al*., 2021), and therefore was applied to wild-type and *tt8-2bp* seeds to measure the baseline germination rates. We sterilized seeds from five different plants with 70% ethanol, 1.5-hour treatment at 6.8% chlorine gas concentration, and 4-hour treatment at 6.8% chlorine gas concentration to test whether chlorine gas treatment reduced the germination rates of *tt8-2bp* more than wild-type seeds (Materials and Methods). No germinated seeds were observed on day 1 after germination started (**Fig. 9B**). We performed two-way ANOVA tests to determine on each day after germination started whether the germination rates of wild-type *vs*. *tt8-2bp* seeds were significantly different, whether the germination rates after 1.5 *vs*. 4 hours of chlorine gas treatment were different, and whether the germination rates of *tt8-2bp* were affected more than wild type when longer chlorine gas treatment was applied (Materials and Methods). The significance threshold was p-value ≤0.05. The statistical test results suggest that significant differences in germination rates were observed between the two different genotypes and between the two different treatment times on day 2 and day 3 after germination started (**Fig. 9B**, **Tab. 2** and **3**).

**Table 2.** P-values of two-way ANOVA tests for identifying differences in germination rates of wild-type and *tt8-2bp* seeds after 1.5 and 4 hours of chlorine gas treatment, respectively. The statistical test is performed for germination rates on each day after germination started. The significance threshold is p≤0.05. The significant p-values are in red letters.

|  | d1 | d2 | d3 | d4 | d5 | d6 | d7 | d8 |
| --- | --- | --- | --- | --- | --- | --- | --- | --- |
| Effect of genotype (G) | NA | 0.002 | 0.005 | 0.083 | 0.423 | 0.73 | 0.2 | 0.14 |
| Effect of treatment time (T) | NA | 0.006 | 0.004 | 0.278 | 0.77 | 0.38 | 0.49 | 0.52 |
| Effect of interaction (GxT) | NA | 0.048 | 0.331 | 0.32 | 0.97 | 0.93 | 0.49 | 0.52 |

**Table 3.** F-statistics of two-way ANOVA tests for identifying differences in germination rates of wild-type and *tt8-2bp* seeds after 1.5 and 4 hours of chlorine gas treatment, respectively. The statistical test is performed for germination rates on each day after germination started.

|  | d1 | d2 | d3 | d4 | d5 | d6 | d7 | d8 |
| --- | --- | --- | --- | --- | --- | --- | --- | --- |
| Effect of genotype (G) | NA | 21.62 | 14.99 | 3.93 | 0.713 | 0.13 | 1.92 | 2.69 |
| Effect of treatment time (T) | NA | 14.12 | 15.34 | 1.37 | 0.095 | 0.86 | 0.53 | 0.46 |
| Effect of interaction (GxT) | NA | 5.41 | 1.07 | 1.1 | 0.001 | 0.008 | 0.53 | 0.46 |

Additionally, p-value of the interaction effect (genotype x treatment time) on germination rates was 0.048 on day 2 (**Tab. 2** and **3**), suggesting that on day 2 the effect of chlorine gas treatment time on germination rates was significantly affected by genotypes. On day 2 and day 3, *tt8-2bp* seeds had lower germination rates than wild-type seeds after 1.5 and 4 hours of chlorine gas treatment, and seed germination rates were lower for both genotypes after longer chlorine gas treatment (**Fig. 9B**). For germination rates on day 4 to 8, no significant differences were found (**Fig. 9B**, **Tab. 2** and **3**). The effect of sterilization with 70% ethanol on seed germination rates of wild type *vs*. tt8-2bp was similar to that of chlorine gas. Based on two-tailed t-tests, *tt8-2*bp had significantly lower germination rates than wild-type seeds on day 3 and day 4 after germination started, but no difference in germination rates were found on day 5-8 (p-value ≤0.05, **Tab. 4**, **Supplementary Fig. S9**). Moreover, there was no significant difference in seed germination rates of wild-type and *tt8-2bp*, respectively, on day 8 after surface sterilization with 70% ethanol, 1.5-hour chlorine gas, and 4-hour chlorine gas (**Fig. 9C**). In summary, under our test conditions, we did not find significant decrease in final *tt8-2bp* seed germination rates compared to wild type after eight days of germination period, suggesting that there was no significant difference in *tt8-2bp vs*. wild-type seed coat permeability toward chlorine gas.

**Table 4.**
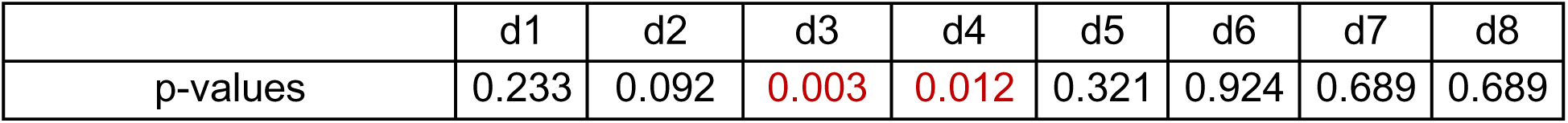
P-values of two-tailed student’s t-test on germination rates of seed sterilized by 70% ethanol: The statistical test is performed for germination rates on each day after germination started. The significance threshold is p ≤ 0.05. The significant p-values are in red letters.

## Discussion

In this study, we performed in-depth anatomical and histochemical characterization of developing pennycress seeds at 7-27 DAP, and did not detect significant changes in the development rate or tissue anatomy of *tt8-2bp* embryo and endosperm compared to those of wild-type seeds. Similar to Arabidopsis *tt8* KO mutant (Chen *et al*., 2014), we found drastic changes in the anatomy and chemical compositions of pennycress *tt8-2bp* seed coats caused by PA deficiency. Based on spatial metabolomics, in addition to complete loss of PAs, 27 DAP *tt8-2bp* seeds had increased disaccharides, trisaccharides, several fatty acids commonly found in seed oils, and amino acid derivatives including glucosinolate and glutathione. Because the thickened oi1 cell wall eventually became the hardened coat protecting endosperm and embryo from the outer environment, we analyzed the chemical compositions of this outer seed coat of mature wild-type and *tt8-2bp* seeds *via* ss NMR. In wild-type outer seed coats, PAs were the main phenolic polymers, and fully hydrated mature outer seed coats of *tt8-2bp* had decreased PAs and cell wall polysaccharides. Consistently, mature *tt8-2bp* seeds had decreased seed coat dry weights and increased embryo dry weights. Additionally, PA deficiency in *tt8-2bp* seed coats also correlated with increased seed coat permeability toward water-soluble molecules and imbibition rates.

### Seed coat anatomy and chemical compositions

Seed development and anatomy are well-characterized in Arabidopsis and *B. napus* (Wan *et al*., 2002; Haughn and Chaudhury, 2005; Gao *et al*., 2022a). The embryo and endosperm development of Spring32 pennycress underwent similar anatomical and morphological changes as Arabidopsis and *B. napus* (**Fig. 2**). However, there are several differences in the seed coat anatomy of pennycress, Arabidopsis and *B. napus.* Pennycress seed coats have three ii and oi layers (**Fig. 3A**) whereas Arabidopsis seed coats have three ii layers and two oi layers (Haughn and Chaudhury, 2005; Huang *et al*., 2023). *B. napus* was reported to have 6-8 ii layers at flowering, 2-3 ii layers when the embryo was at torpedo stage, and four layers of oi throughout seed development (Wan *et al*., 2002). Regardless of the variations in ii and oi cell layers, seeds of all three species accumulated PAs in ii1 and formed thickened cell walls outside oi1 (**Fig. 3B, 4A**; Windsor *et al*., 2000; Haughn and Chaudhury, 2005; Zhang *et al*., 2022), suggesting that the PA accumulation in ii1 and formation of thickened oi1 cell wall may be evolutionally conserved. Interestingly, although seeds of pennycress, Arabidopsis and *B. napus* all develop thickened secondary cell walls in oi1 layer, the cell wall chemical compositions are different. Pennycress wild-type seeds are the only ones accumulating PAs in the thickened oi1 cell wall (**Fig. 3B**, **5B**, **6A**), whereas no PAs are detected in the thickened oi1 cell wall of Arabidopsis (Windsor *et al*., 2000) or *B. napus* (Gao *et al*., 2022b). Although one *B. napus* study claimed to have detected PAs in the seed coat palisade layer (*i.e.*, oi1 cells), the histology evidence was shown in low-magnification and low-resolution micrographs where different integument layers cannot be clearly identified and thus cannot sufficiently support the claim (Qu *et al*., 2013). Additionally, among the three species, only Arabidopsis seed coats produce mucilage in the oi2 cells (Jaimie *et al*., 2005; Haughn and Western, 2012), and pennycress seeds did not have mucilage that extruded to cover the entire seed (**Supplemental Fig. S6B**).

Seed coat lignin and PAs are large polymers of phenolic compounds and main sources of seed fibers. When quantifying seed fiber contents, lignin and PAs are usually not distinguished (Steck *et al*., 2022). However, from the perspective of seed quality and nutrient profile improvement, it is important to know the proportions of lignin and PAs in the total seed fibers so specific metabolic pathways can be targeted. When lignin and cell wall-associated PAs in the outer seed coats were quantified separately, seeds of different plants were found to have different proportions of lignin and PAs in their outer seed coats (Steck *et al*., 2022). For example, grape outer seed coats have both lignin and PAs, but chokeberry seed outer seed coats only had PAs whereas raspberry outer seed coats only had lignin (Steck *et al*., 2022). In pennycress wild-type and *tt8-2bp* seeds, lignin was accumulated in small regions of the thickened oi1 cell wall (**Fig. 4C**) whereas in wild-type seeds PAs were integrated into the entire thickened oi1 cell wall and turned the cell wall brown after PA oxidation to condensed tannins (**Fig. 4B**, **Supplemental Fig. S2**). Based on the ss NMR analysis, the PA-deficient outer seed coats of *tt8-2bp* had barely detectable phenolic compounds compared to wild-type outer seed coats (**Fig. 6A**), suggesting that PAs were the main phenolic compounds and fiber sources in wild-type, dark-colored pennycress seeds. Therefore, to reduce seed fiber contents of wild-type pennycress seeds, targeting PA accumulation should be more effective than targeting lignin biosynthesis and polymerization pathways, supporting *TT2* and *TT8* as good gene targets for pennycress domestication.

*tt8-2bp* seeds had increased seed coat permeability to water soluble molecules and imbibition rates (**Fig. 8A**, **9A**), and it can be attributed to PA deficiency in ii1 layers. However, this may not be the only cause of increased imbibition rates and seed coat permeability. In Arabidopsis, in addition to PA deficiency, *tt* KO mutant seeds had defects in a cuticle layer formed by ii1 cells and showed increased seed coat permeability to water soluble molecules (DeBolt *et al*., 2009; Loubery *et al*., 2018).

There could be similar cuticle defects in pennycress *tt8-2bp* seeds and the cause of increased *tt8-2bp* seed coat permeability needs to be investigated further before a conclusion can be reached. The increased imbibition rates of *tt8* KO seeds can lead to a faster germination in the field when the soil moisture level and temperature are favorable. The increased seed coat permeability of *tt8* KO can affect seed germination rates if the seeds need to be pre-treated with chemicals before planting. If *tt8* KO mutant seeds are soaked in chemicals such as fungicides for too long that the fungicides permeate across seed coats and damage embryos, it can lead to reduced seed germination in the field. Any potential negative effects could be avoided by first testing the treatment on a small batch of seeds to optimize the treatment condition.

### Altered nutrient partitioning between the embryo and seed coat of *tt* mutants

In correlation with PA deficiency in *tt8-2bp* seed coats, spatial metabolomic analysis on wild-type and mutant seeds at 27 DAP showed that *tt8-2bp* seeds had increased small sugars such as sucrose, raffinose, *etc*. (**Fig. 5D-E**, **Supplementary Fig. S7D-E**) and fatty acids such as linoleic acid, oleic acid, palmitic acid, and α-Linolenic acid (**Fig. 5G**, **Supplementary Fig. S7O-T**). The small sugars and fatty acids are main substrates in starch and storage lipid synthesis in embryos, suggesting that *tt8-2bp* mature seeds may have more storage nutrients fixed in the embryo than wild-type seeds. Indeed, mature *tt8-2bp* seeds had higher embryo dry weights and lower seed coat dry weights than wild-type seeds, which led to a 9% increase in the total nutrients measured by dry weights in *tt8-2bp* mature embryos (**Fig. 7C**, **7D**). Consistently, *tt8-2bp* seeds had ∼10% increase in total seed oil compared to wild-type seeds. Chen *et al*. (2014) found evidence supporting *TT8*’s role as a negative regulator of fatty acid biosynthesis in Arabidopsis. Increased seed storage nutrients were also observed in *tt* KO mutants of other *Brassicaceae* plants. Arabidopsis *ttg1* KO mutant seeds had increased embryo dry weights compared to wild-type seeds which was also accompanied by an increase in total starch, proteins, and fatty acids (Chen *et al*., 2015b). In *Camelina sativa* L. and *B. napus*, disruption of *TT* genes led to increase in total fatty acids and storage lipids of mature seeds (Zhai *et al*., 2020; Cai *et al*., 2024; Li *et al*., 2024). These results indicate that the function of *TT* genes in controlling seed coat PA synthesis and modulating seed storage nutrient accumulation especially fatty acids and lipids are likely conserved in *Brassicaceae* family.

The increase in fatty acid and oil contents of *tt* mutant seeds is highly correlated with altered nutrient partitioning between seed coats and embryos. Positive correlations were found between seed oil content *vs*. sucrose, starch, and storage proteins in embryos (Chen *et al*., 2015b; Zhai *et al*., 2020; Apriyanto *et al*., 2022). In contrast, significant negative correlations were found between seed oil content *vs*. seed coat thickness and cell wall components such as lignin, cellulose, and hemicellulose (Miao *et al*., 2019; Zhang *et al*., 2022). Consistently, a yellow-seed pennycress mutant with increased seed oil content was shown to have decreased seed coat thickness (Griffiths *et al*., 2025). In *B. napus*, *TT8* expression had a positive correlation with seed coat contents (seed coat weight to total seed weight ratios) and a negative correlation with seed oil content (Zhang *et al*., 2022). In Arabidopsis and *B. napus*, *ttg1, tt2,* and *tt8* KO mutations led to down-regulation of key genes in flavonoid biosynthesis and up-regulation of key genes in fatty acid biosynthesis (Chen *et al*., 2012; Chen *et al*., 2014; Chen *et al*., 2015b; Li *et al*., 2024). In summary, all three members of the *TTG1-TT2-TT8* transcription factor complex regulate gene expression of key genes involved in PA biosynthesis and fatty acid biosynthesis and in turn, modulate nutrient partitioning between seed coats and embryos. Therefore, *TTG1*, *TT2*, and *TT8* are good targets for improving seed quality and nutrient contents of pennycress and other oil crops in the *Brassicaceae* family.

In this study, we showed that a KO mutation in pennycress *TT8* led to reduced seed coat fiber and dry weights and increased seed fatty acids and embryo dry weights without anatomical defects in embryo or endosperm. The increased seed imbibition rates of *tt8-2bp* seeds could be beneficial for faster seed germination in the field, but the increased *tt8-2bp* seed coat permeability to water soluble molecules warrant more testing if chemical treatments of seeds were to be applied before planting. Overall, we did not find agronomically unfavorable traits in *tt8-2bp* seeds. Further field studies on *tt8-2bp* seed germination rates under abiotic stresses are needed to evaluate its efficiency of seedling establishment under drought stress, salt stress, or water-logging condition.

## Supplementary data

Table S1. Incubation steps for seed sample dehydration and paraffin embedding.

Table S2. Incubation steps for seed paraffin section de-paraffin and rehydration before histological staining.

Figure S1. Timeline and representative images of Spring32 wild-type and *tt8-2bp* plant development from seedling establishment to seed maturity.

Figure S2. Pictures of developing and mature seeds and embryos of Spring32 wild-type and *tt8-2bp*, respectively.

Figure S3. Main seed tissues of a developing Spring32 wild-type pennycress seed.

Figure S4. Additional micrographs of safranin O and alcian blue stained wild-type and *tt8-2bp* seed paraffin sections at 7-27 DAP.

Figure S5. Paraffin sections of 15 DAP wild-type and *tt8-2bp* seeds unstained and stained for detection of PAs.

Figure S6. The thickened oi1 cell wall of wild-type and *tt8-2bp* were stained differently by alcian blue at 15-27 DAP and detection of mucilage and pectin in pennycress seeds.

Figure S7. Additional molecular features identified in the spatial metabolomics of small molecules and lipids in wild-type and *tt8-2bp* seeds at 27 DAP.

Figure S8. Ss NMR spectrum of pure potassium bromide showing lack of background noise in ^13^C spectra of seed coats.

Figure S9. Boxplots of wild-type and *tt8-2bp* seed germination rates after seed surface sterilization with 70% ethanol.

## Acknowledgements

We would like to thank Mr. Will Chrisler and Ms. Marija Velickovic at EMSL, PNNL (WA., US) for their support and advice on histological analyses and microscopy done in this study. We would like to also thank Ms. Tanya Winkler at EMSL, PNNL (WA., US) for her support and advice on plant caretaking and locating instruments used for this study.

## Author Contribution

XD, JCS, and PH: conceptualization; XD, AL, and DV: methodology; XD, AL, and DV: formal analysis; XD, SD, MS, AL, and DV: investigation; BG and JCS: resources; XD, AL, and DV: data curation; XD and AL: writing - original draft; XD, SD, BG, AL, DV, JCS, and PH: writing - review & editing; XD, SD, and AL: visualization; PH: supervision; JCS and PH: funding acquisition

## Data availability

All the MALDI-MSI images and corresponding brightfield images can be found *via* the following link: https://metaspace2020.org/api_auth/review?prj=6c81d8ac-eb19-11ef-a048-f7b2854b75cc&token=3FuQkohEOr7H

## Abbreviations

*tt8*: *transparent testa 8*
KO: knockout
PAs: proanthocyanidins
ii: inner integument
oi: outer integument
DAP: day after pollination
ss NMR: solid-state nuclear magnetic resonance
MALDI-MSI: matrix-assisted laser desorption ionization mass spectrometry imaging
HCl: hydrochloric acid

