## Supplemental files for "A pennycress *transparent testa 8* knockout mutant has drastic changes in seed coat anatomy and chemical compositions"

Supplementary Table S1. Consecutive incubation steps for seed sample dehydration, clearing, and paraffin embedding. Seed samples were placed in plastic embedding cassettes and the incubation was done in glass beakers covered with alumina foil.

| | **Solution** | **Time** |
|---|---|---|
| Sample dehydration | 60% ethanol | 1 hour |
| | 80% ethanol | 1 hour |
| | 95% ethanol | 1 hour |
| | 100% ethanol | 1 hour |
| | 100% ethanol | Overnight |
| Sample clearing and embedding | 100% xylenes | 1 hour |
| | 100% xylenes | 1.5 hour |
| | Melted paraffin (56-60 ºC) | 1 hour |
| | Melted paraffin (56-60 ºC) | 2 hour |

Supplementary Table S2: Consecutive incubation steps for seed paraffin section deparaffinization and rehydration before histological staining. The paraffin sections were rehydrated to the same water content as the staining solution to be used.

| **Solution** | **Time** |
|---|---|
| 100% xylenes | 5 minutes |
| 100% xylenes | 5 minutes |
| 100% ethanol | 3 minutes |
| 100% ethanol | 3 minutes |
| 95% ethanol | 3 minutes |
| 70% ethanol | 3 minutes |
| 50% ethanol | 3 minutes |
| Deionized water | 3 minutes |

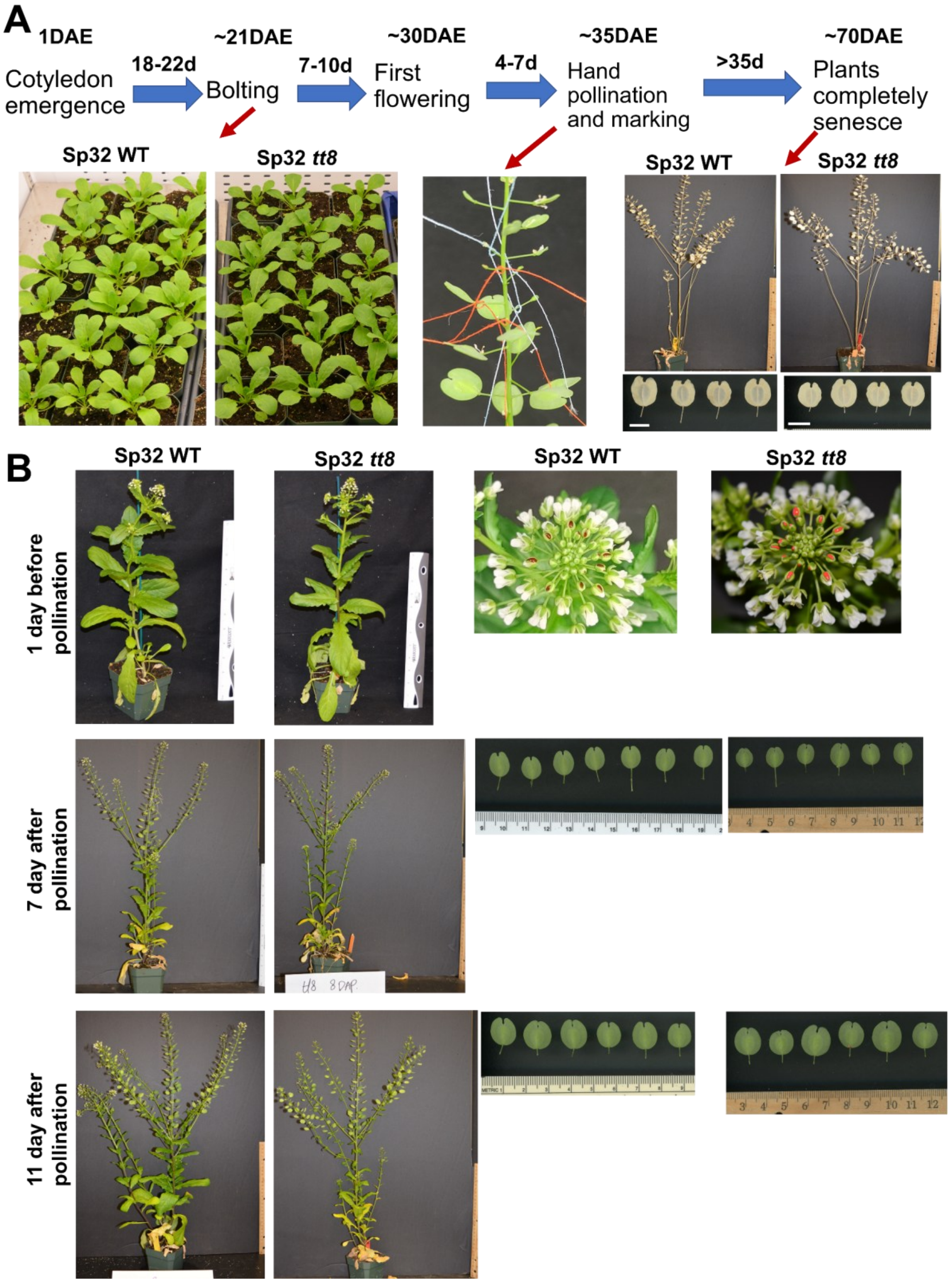
A
1DAE
Cotyledon emergence
18-22d
~21DAE
Bolting
7-10d
~30DAE
First flowering
4-7d
~35DAE
Hand pollination and marking
>35d
~70DAE
Plants completely senesce
Sp32 WT
Sp32 tt8
Sp32 WT
Sp32 tt8
B
Sp32 WT
Sp32 tt8
Sp32 WT
Sp32 tt8
1 day before pollination
7 day after pollination
11 day after pollination
tt8 8 DAP

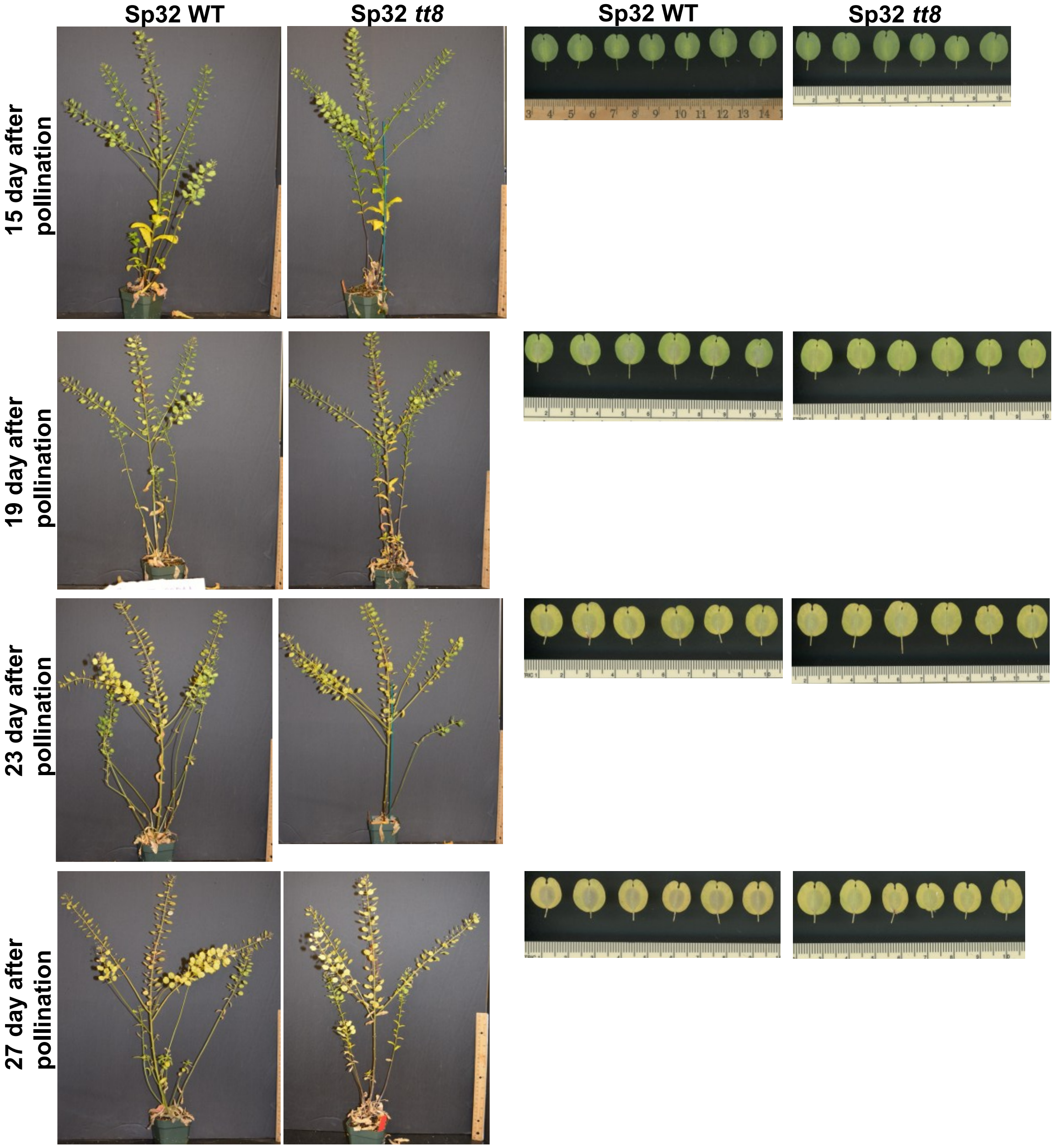


Supplementary Figure S1. Timeline of Spring32 wild-type (WT) and *tt8-2bp* plant development in the growth chamber. (A) Sterilized seeds were stratified for 2 days and germinated on filter paper. Germinated seedlings were planted and grown at 22 ºC, 50% humidity, under 200 μmol $m^{-2}$ $s^{-1}$ light. Fertilizer (Jack's Pro) was applied twice a week. Senesced plant pictures were taken on 41 day after pollination (DAP). DAE, day after emergence. Scale bars = 1cm. (B) Representative images of Sp32 WT plants and seed pods at different days after pollination when seed samples were harvested.

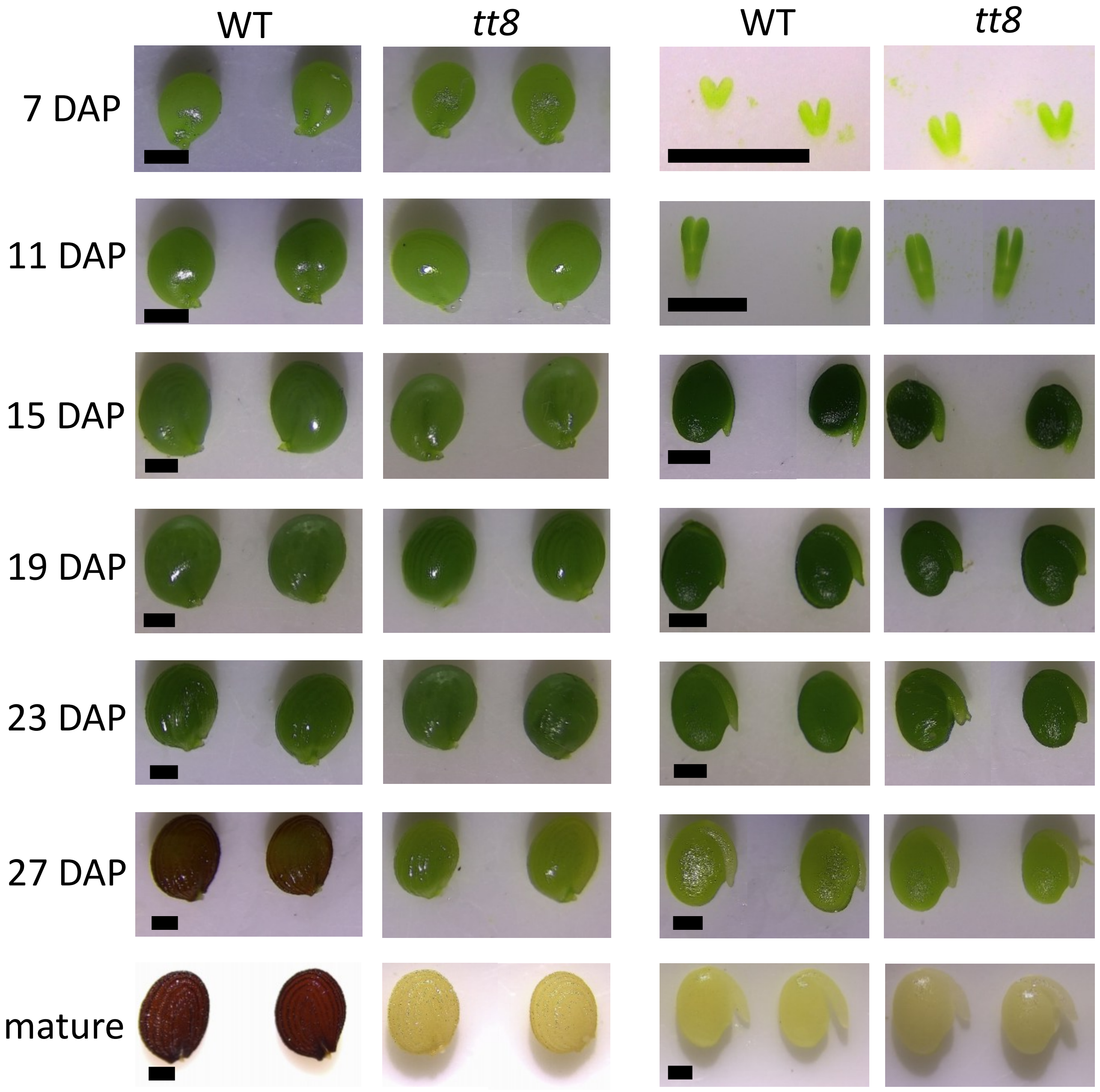


Supplementary Figure S2: Pictures of developing and mature seeds and embryos of Spring32 wild-type (WT) and *tt8-2bp*, respectively. The scale bars are 500 µm.

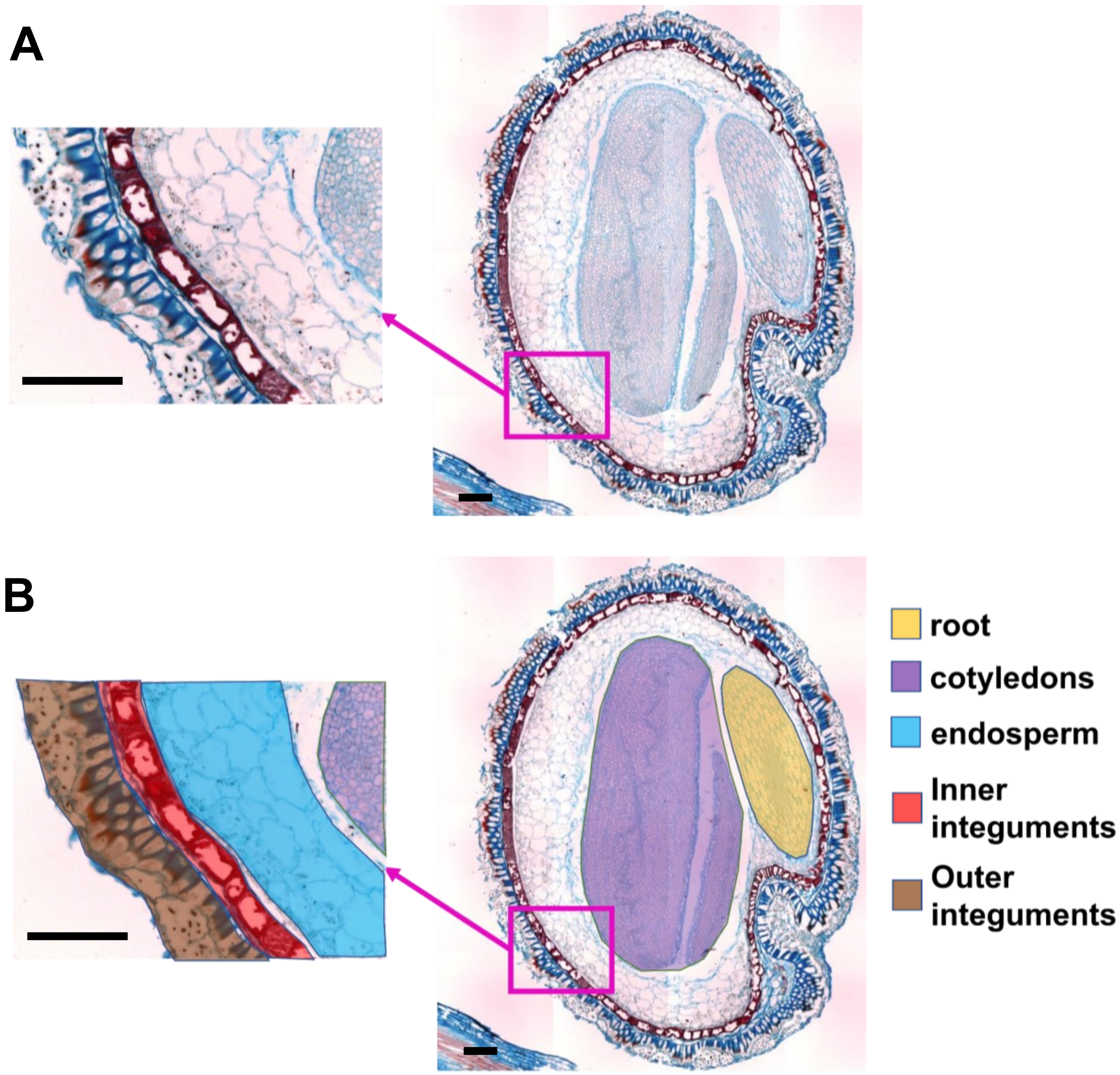


Supplementary Figure S3: Main seed tissues of a developing Spring32 wild-type pennycress seed. (A, B) A seed paraffin section of a WT seed at 15 DAP stained with safranin O and alcian blue and a blowup image focused on the seed coat cell layers. Scale bars = 100 μm.

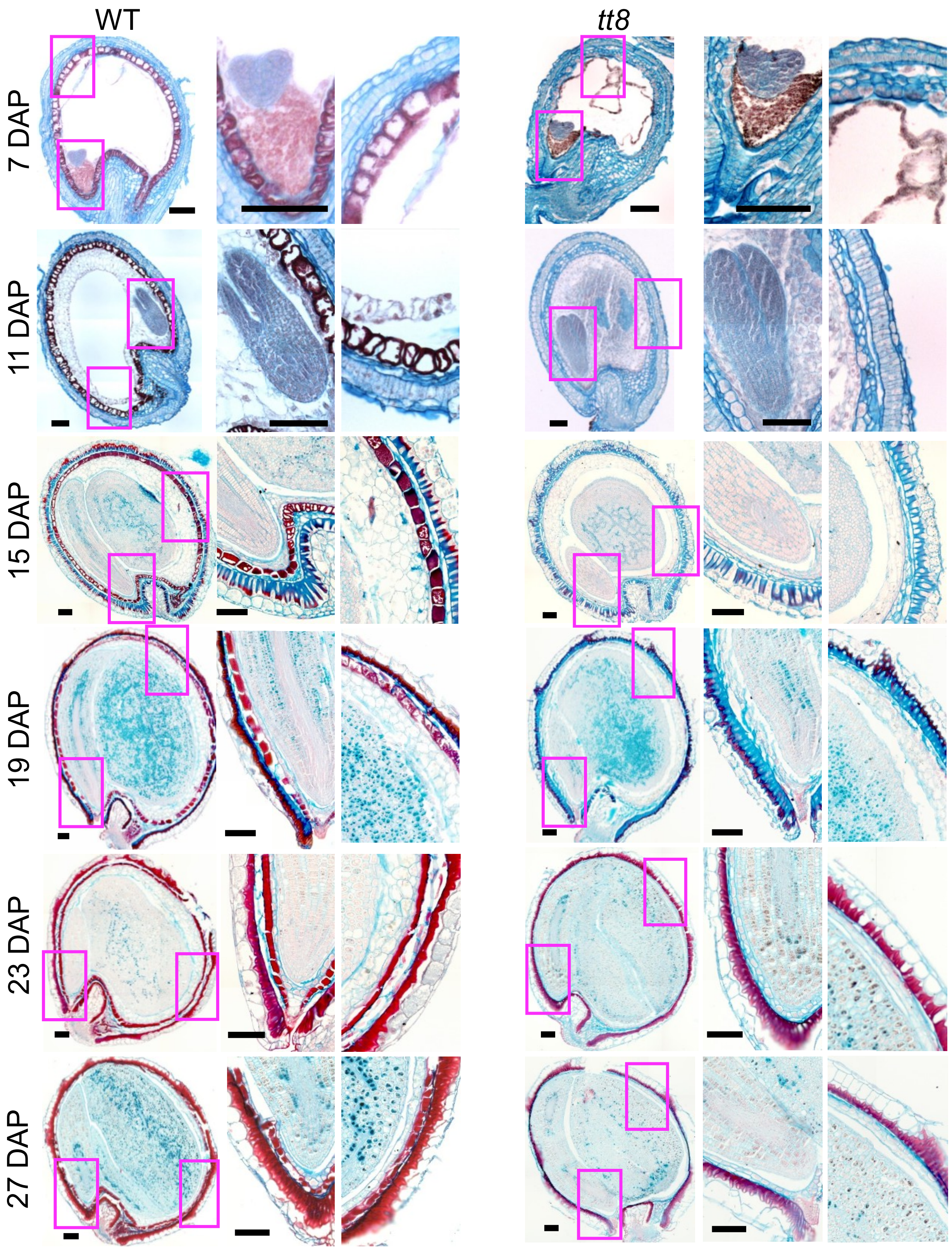


Supplementary Figure S4. Additional micrographs of histological staining of wild-type (WT) and *tt8-2bp* embryo and endosperm shows that the two seed tissues developed at a similar rate at 7-27 DAP. Seed sections were stained with safranin-O (stains secondary cell wall and nuclei red) and counter-stained with alcian blue (stains acidic polysaccharides of primary cell wall blue). Blowup images of shown on the right of each seed micrograph. Scale bars = 100 µm.

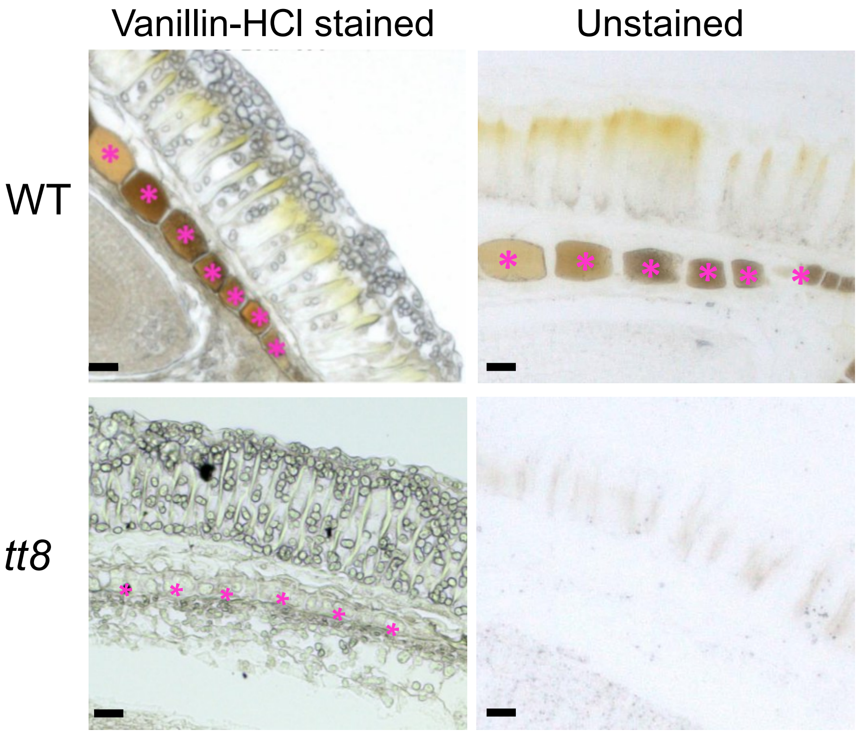


Supplementary Figure S5. Paraffin sections of 15 DAP wild-type (WT) and *tt8-2bp* seeds unstained and stained with vanillin-HCl for detection of PAs. Magenta asterisks mark the ii1 cells accumulating PAs. Note that in both WT and *tt8-2bp* sections, there was no difference in the color of the ii1 cells in the unstained *vs.* vanillin-HCl stained seed sections. Scale bars = 25 μm.

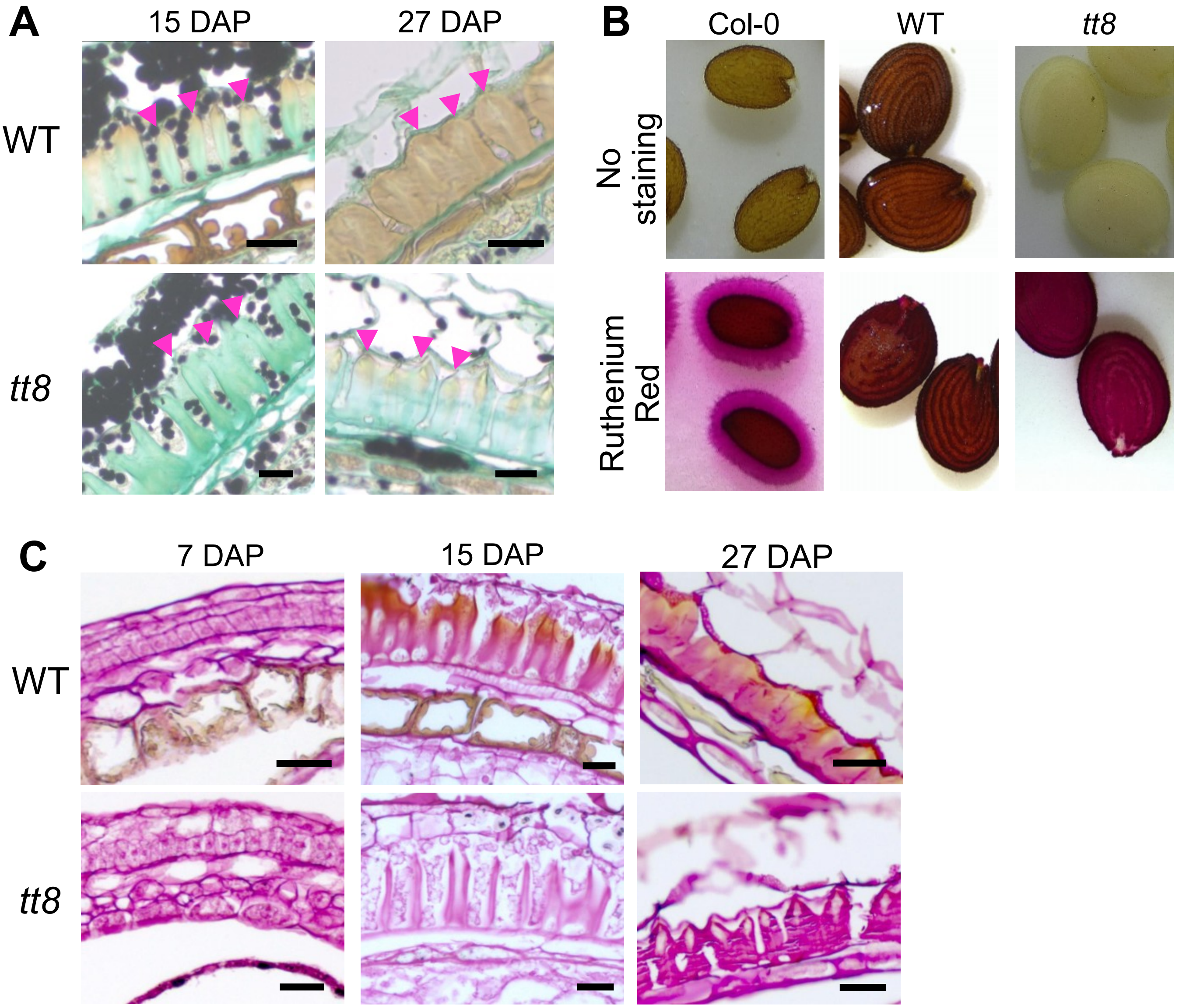


Supplementary Figure S6: The thickened oi1 cell wall of wild-type (WT) and *tt8-2bp* were stained differently by alcian blue at 15-27 DAP and detection of mucilage and pectin in pennycress seeds. (A) Seed sections of Sp32 WT and *tt8* at 15 and 27 DAP were stained with Alcian blue and Lugol's iodine which stain primary cell wall light turquoise and starch granules black, respectively. (B) Mature seeds of Arabidopsis Col-0, pennycress Sp32 WT, and *tt8* with and without 0.1% ruthenium red staining after seed imbibition in 100mM EDTA (pH 8) to detect secretion of seed mucilage. (C) Pennycress seed paraffin sections stained with 0.1% ruthenium red. Scale bars = 25 μm.

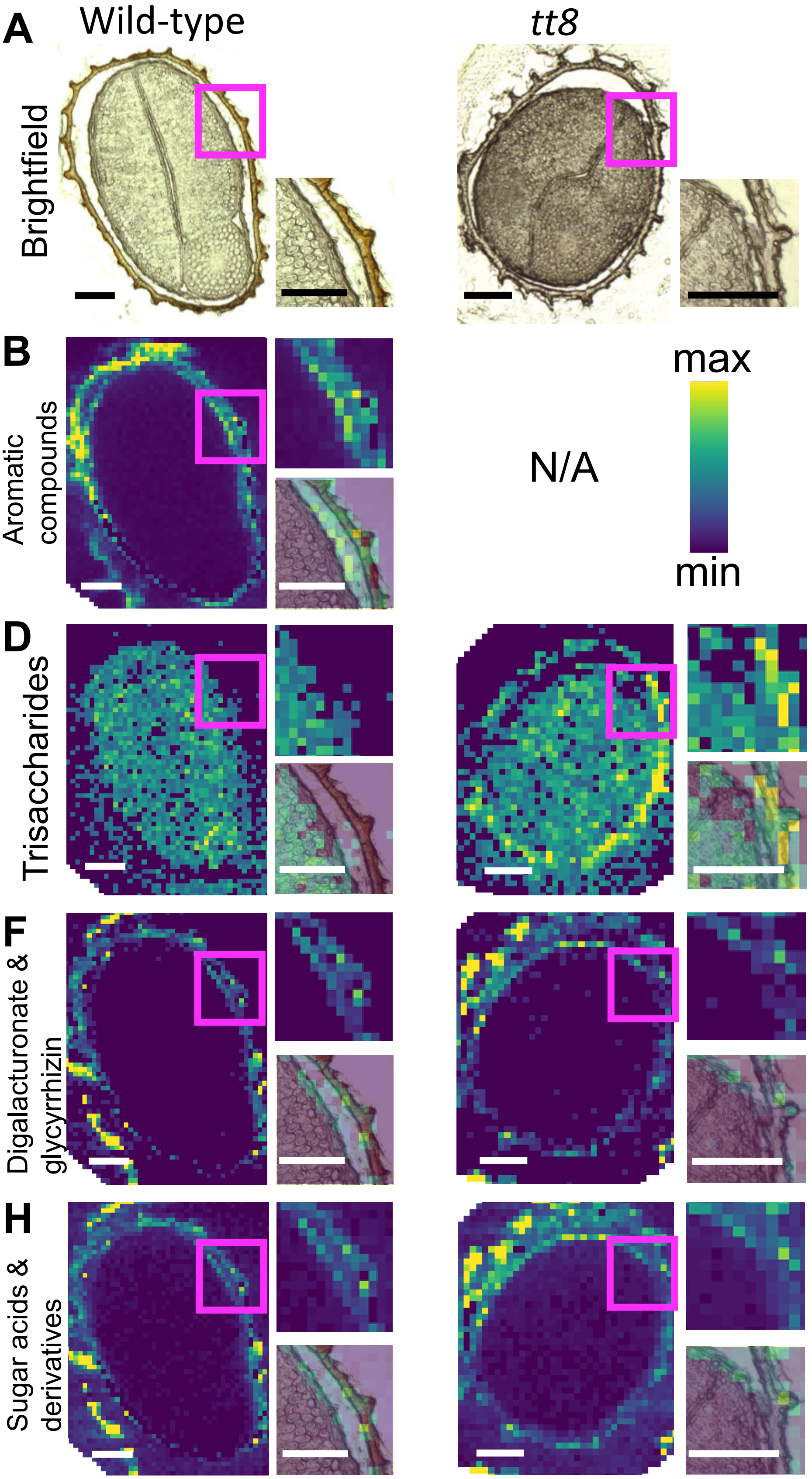

A
Wild-type
tt8
Brightfield
B
Aromatic compounds
N/A
max
min
C
p<0.0001
log2(signal intensity)
25
20
15
10
5
0
WT
tt8
D
Trisaccharides
E
p=0.004
log2(signal intensity)
17.5
17.0
16.5
16.0
WT
tt8
F
Digalacturonate & glycyrrhizin
G
N.S
log2(signal intensity)
18.0
17.5
17.0
16.5
WT
tt8
H
Sugar acids & derivatives
I
N.S
log2(signal intensity)
20.0
19.5
19.0
18.5
WT
tt8

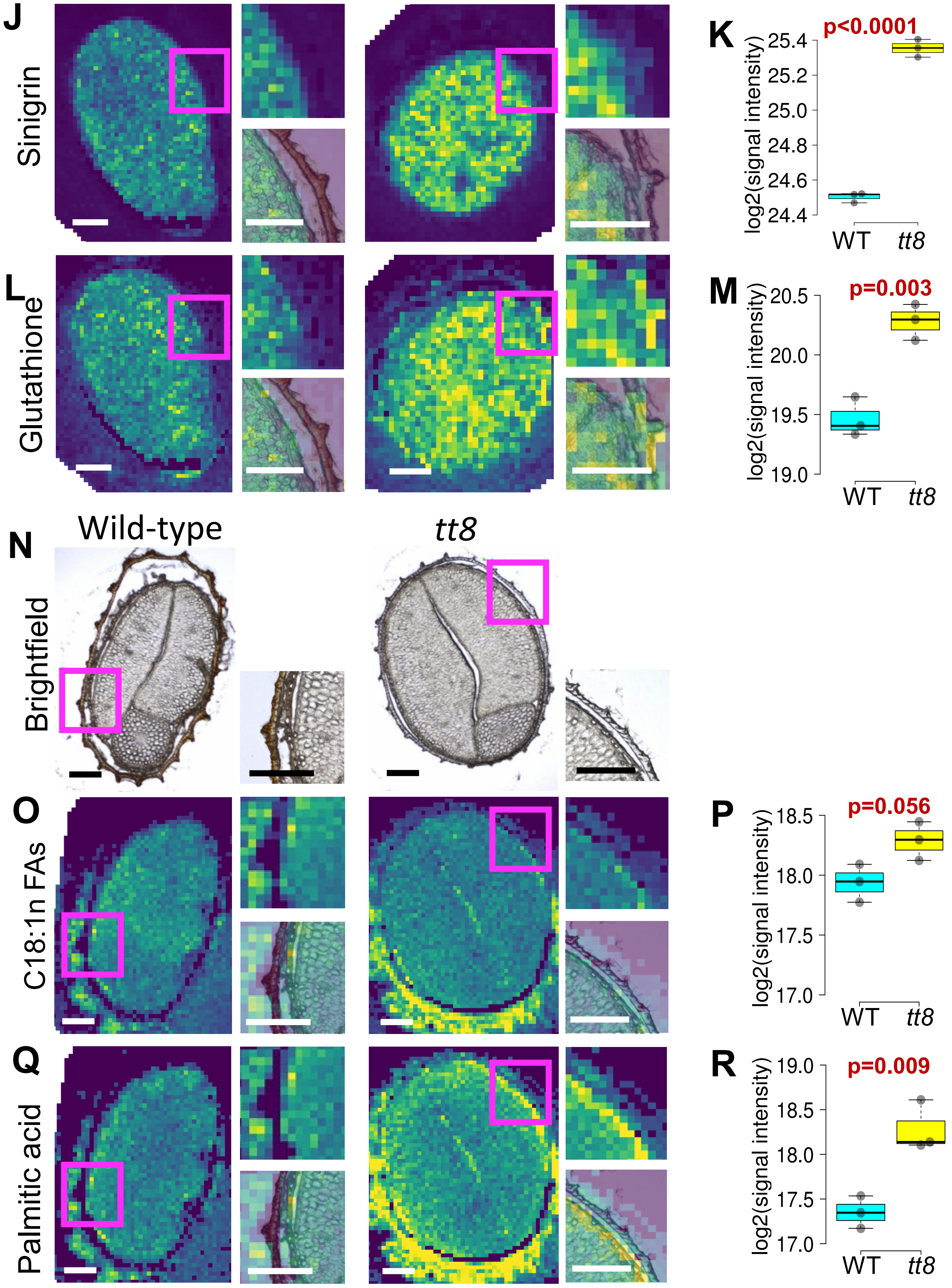

J
Sinigrin
K
p<0.0001
log2(signal intensity)
25.4
25.2
25.0
24.8
24.6
24.4
WT
tt8
L
Glutathione
M
p=0.003
log2(signal intensity)
20.5
20.0
19.5
19.0
WT
tt8
N
Wild-type
tt8
Brightfield
O
C18:1n FAs
P
p=0.056
log2(signal intensity)
18.5
18.0
17.5
17.0
WT
tt8
Q
Palmitic acid
R
p=0.009
log2(signal intensity)
19.0
18.5
18.0
17.5
17.0
WT
tt8

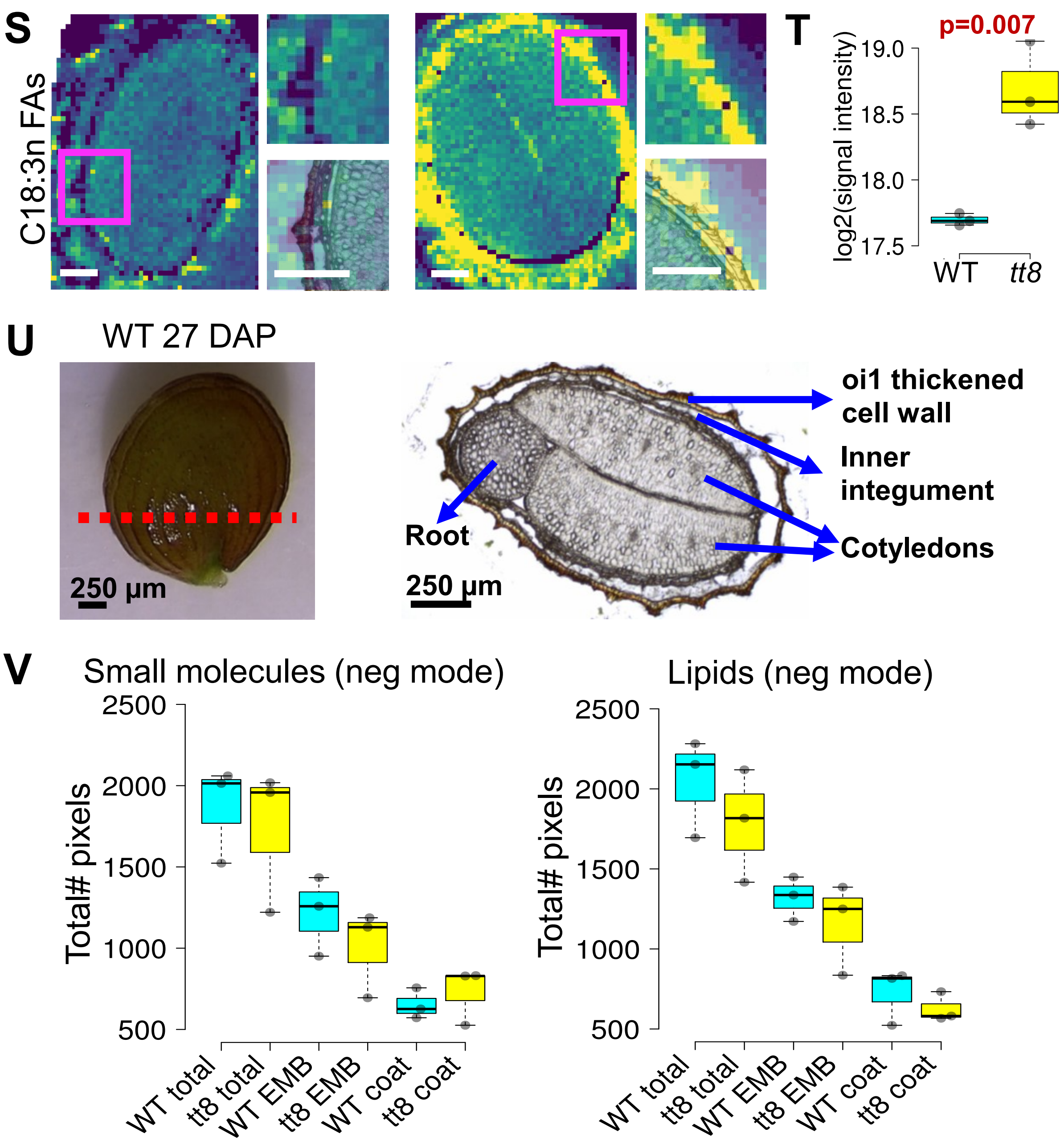

S
C18:3n FAs
T
p=0.007
log2(signal intensity)
19.0
18.5
18.0
17.5
WT
tt8
U
WT 27 DAP
250 µm
oi1 thickened cell wall
Inner integument
Root
Cotyledons
250 µm
V
Small molecules (neg mode)
Lipids (neg mode)
Total# pixels
2500
2000
1500
1000
500
WT total
tt8 total
WT EMB
tt8 EMB
WT coat
tt8 coat

Supplementary Figure S7. Additional molecular features of interests identified in the spatial metabolomic analysis of small molecules and lipids in Spring32 wild-type (WT) and *tt8-2bp* seeds at 27 DAP. (A, N) Representative images of brightfield micrographs of seed cryosections used for MALDI-MSI for the detection of small molecules and lipids, respectively, under the negative ion mode. A blowup image of the indicated seed region (magenta box) is provided to show the spatial separation of seed coat and embryo. (B, D, F, H, J, L, O, Q, S) Heatmaps showing the ion signal intensities of some small molecules and fatty acids detected in the same seed sections show in panel A and H. The heatmaps of the three different molecular features are displayed in the same color scale with deep blue representing the min value and yellow representing the max value without any data transformation. The min and max values are the same in the heatmaps of wild-type and *tt8* seed sections of the same molecular feature. A blowup image of the indicated seed region (magenta box) is shown to the upper right of each heatmap, and an overlaid image of the blowup heatmap adjusted to 50% transparency and the brightfield micrograph is shown to the lower right of each map. Scale bars = 200 μm. The small molecules include aromatic compounds ($C_{15}H_{12}O_6$), trisaccharides ($C_{18}H_{32}O_{16}$), digalacturonate & glycyrrhizin ($C_{12}H_{18}O_{13}$), sugar acids & derivatives ($C_6H_{10}O_7$), sinigrin ($C_{10}H_{17}NO_9S_2$), and glutathione ($C_{10}H_{17}N_3O_6S$). The fatty acids include C18:1n fatty acids (C18:1n FAs, $C_{18}H_{34}O_2$), palmitic acid ($C_{16}H_{32}O_2$), and C18:3n fatty acids (C18:3n, $C_{18}H_{30}O_2$). There are 14 different aromatic compounds identified for $C_{15}H_{12}O_6$ with m/z of 287.0561, of which nine are flavonoids such as dihydrokaempferol, fustin, etc. There are 14 different trisaccharides such as raffinose, maltotriose, etc., identified for $C_{18}H_{32}O_{16}$ with m/z of 503.1618. Digalacturonate and glycyrrhizin are identified for $C_{12}H_{18}O_{13}$ with m/z of 369.0675. There are 20 different sugar acids and sugar acid derivatives such as D-galacturonic acid, L-iduronic acid, etc. identified for $C_6H_{10}O_7$ with m/z of 193.0354. Sinigrin is the only metabolite identified for $C_{10}H_{17}NO_9S_2$ with m/z of 358.0272. Glutathione is the only metabolite identified for $C_{10}H_{17}N_3O_6S$ with m/z of 306.0765. There are five different C18:1n fatty acids (C18:1n FAs, $C_{18}H_{34}O_2$) including oleic acid, 18:1(6Z), 18:1(11Z), 18:1(11E), and (9E)-octadecenoate with m/z of 281.2486. Palmitic acid is the only metabolite identified for $C_{16}H_{32}O_2$ with m/z of 255.2330. Two fatty acids including α-Linolenic acid and18:3(6Z,9Z,12Z) are identified for $C_{18}H_{30}O_2$ with m/z of 277.2173. (C, E, G, I, K, M, P, R, T) Boxplots of log2-transformed average ion signal intensities of each molecular feature detected in three wild-type and *tt8* seed sections, respectively. The p-values calculated by two-tailed student's t-tests for comparing the log2-transformed average ion signal intensities between wild-type and *tt8* seed sections are shown above the boxplots. The gray dots represent values of different seed sections. N.S. means "not significant" with the significance threshold of 0.05. (U) A photo of a Sp32 wild-type seed at 27 DAP (left) and a brightfield micrograph of a 27 DAP WT seed cryosection without any staining (right). The red dotted line indicate the direction of the plane where seed cryosections were acquired. (V) Boxplots of the total number of pixels in the whole seed (total), embryo (EMB), and seed coat (coat) regions of three wild-type and *tt8-2bp* seed sections, respectively, used for analyzing small molecules and lipids under negative ion mode. The gray dots represent values of different seed sections. Two-tailed student's t-tests were performed to compare the total number of pixels in the same seed compartment between wild-type and *tt8* seed sections. There is no significant difference detected in any comparison with the significance threshold of 0.05.

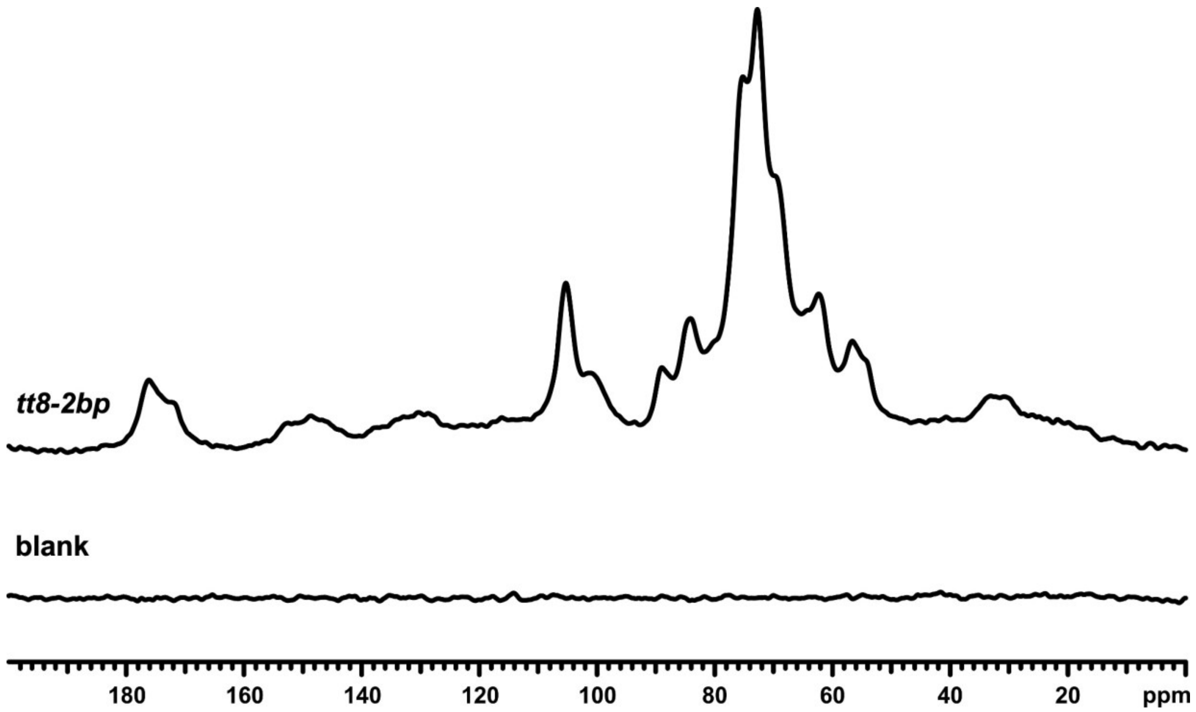


Supplementary Figure S8. The $^{13}C$ CP/MAS spectra of *tt8-2bp* and a rotor full of potassium bromide to serve as a blank. Both spectra were acquired under identical conditions and are plotted on the same scale. Recycle delay was 2s and total number of transients was 65536 for each.

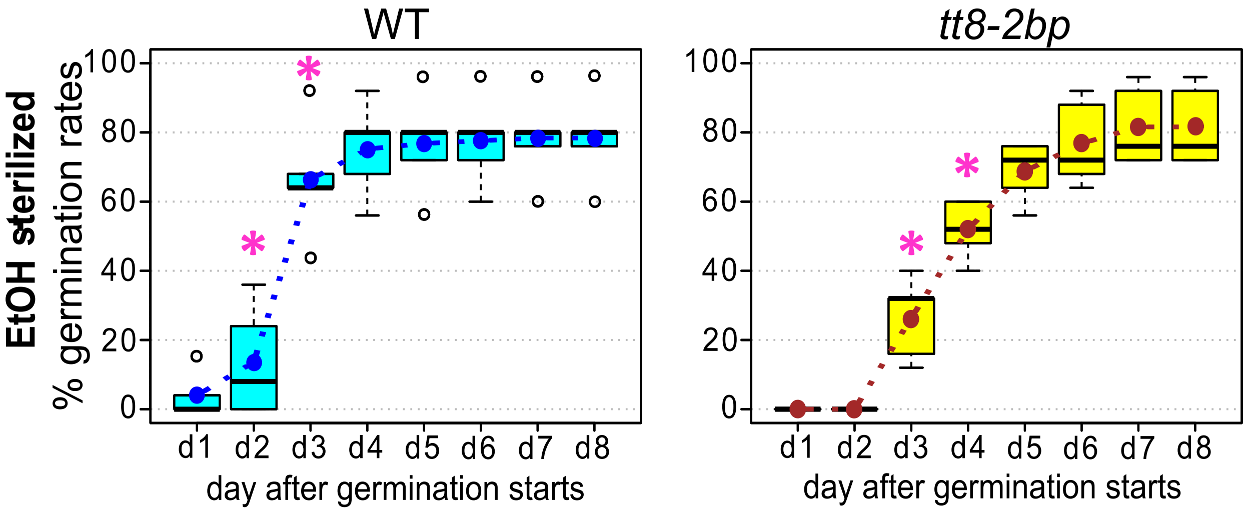


Supplementary Figure S9. Boxplots of wild-type (WT) and *tt8-2bp* seed germination rates after seed surface sterilization with 70% ethanol (EtOH). Asterisks indicate statistically significant difference (p-value ≤0.05) between the germination rates of WT and *tt8-2bp* based on two-tailed student's t-tests.
